# Niche Macrophages Recycle Iron to Tumor Cells and Foster Erythroblast Mimicry to Promote Bone Metastasis and Anemia

**DOI:** 10.1101/2025.04.23.650120

**Authors:** Yujiao Han, Hirak Sarkar, Zhan Xu, Sereno Lopez-Darwin, Yong Wei, Xiang Hang, Fengshuo Liu, Kimberley Tran, Wei Wang, Jennifer M. Miller, Christina J. DeCoste, Dylan Blohm, Robert L. Satcher, Xiang H.-F. Zhang, Yibin Kang

## Abstract

Bone marrow is both a primary site for blood cell production and a fertile niche for metastatic cancer cell growth, notably in breast cancer. Although anemia is common among patients with bone metastasis, the mechanistic link between metastatic colonization and disrupted erythropoiesis remains poorly understood. Using *in vivo* niche labeling and single-cell RNA sequencing, we identified a specialized population of VCAM1^+^CD163^+^CCR3^+^ macrophages enriched in the bone metastatic niche. These macrophages, typically essential for erythropoiesis in healthy bone marrow, are co-opted by tumor cells to support their growth through iron acquisition. The hijacking of these macrophages by tumor cells reduces iron availability for erythroblasts, impairing erythropoiesis and contributing to anemia. With increased iron supply, tumor cells further adapt by mimicking erythroblasts, producing hemoglobin under GATA1 regulation in response to hypoxic stress. Notably, macrophages with similar iron-regulating features were found in human bone metastases across multiple cancer types, and elevated *HBB* expression in breast cancer correlates with increased risk of bone metastasis. These findings establish iron-recycling macrophages as essential regulators within the metastatic bone niche, revealing novel insights into the interplay between immune modulation, metal metabolism and tumor cell plasticity in driving metastatic progression and anemia.

## Introduction

As a vital organ that provides both structural support and blood cell production, bone hosts a biologically active microenvironment that regulates the dynamics of stem, progenitor, and mature cells from various lineages ^1,2^. It also serves as a common site for the metastatic spread of cancer cells ^3–6^. Patients with bone metastasis often suffer severe and life-threatening local and systemic complications, including anemia, leukopenia, thrombocytopenia, hypercalcemia, muscle weakness, spinal cord compression, fracture and bone pain ^4,7^. Bone-resorbing osteoclasts have been a central focus in studies of tumor-stromal interactions in bone metastasis and are the primary therapeutic targets of agents such as bisphosphonates and RANKL-neutralizing antibodies. While these treatments help alleviate complications from bone metastasis, they often do not result in a cure or provide significant survival benefits for cancer patients ^8,9^. In addition to osteoclasts, previous research has identified key interactions between cancer cells and mesenchymal stem cells (MSCs) ^10–12^, osteoblasts ^13–15^, and endothelial cells ^16,17^ during bone metastasis. Hematopoietic stem cells (HSCs), which give rise to most bone marrow cells—including erythrocytes, monocytes/macrophages, granulocytes, platelets, and lymphocytes—undergo tightly regulated processes of production, differentiation, and maturation to maintain homeostasis throughout life ^2,18^.

Hematopoietic cells interact intricately within their lineages and with other stromal cells across spatially distinct regions of bone ^2,18^. However, how metastatic tumor cells interact with hematopoietic cells and the pathological consequences of these interactions remain unclear. An unbiased and comprehensive approach is essential for capturing and profiling the cellular landscape of the bone marrow microenvironment, particularly in regions near the tumor, referred to as the metastatic bone niche.

Disseminated cancer cells face physical, metabolic, and immunologic challenges in distant organs, yet they manage to thrive by accessing essential oxygen and nutrients for survival and proliferation ^19^. Located within the medullary cavities of bones, bone marrow exhibits lower oxygen levels compared to other organs ^2^. This hypoxic condition mirrors the environment in which HSCs originally evolved, providing an adaptive advantage that is essential for maintaining their quiescence and stemness ^18,20,21^.

Tumor cells that infiltrate the HSC niche ^18,22^ must develop mechanisms to survive in this hypoxic environment.

Bone is a nutrient-rich tissue, containing abundant minerals—including calcium, phosphorus, magnesium, iron, and zinc—as well as fat, fat-soluble vitamins, and amino acids ^23^. However, these nutrients are often not directly accessible to disseminated tumor cells, which may hijack or recruit bone stromal cells to obtain access to them for survival and proliferation. Previous studies have shown that tumor-induced osteoclastic bone resorption releases calcium (Ca^2+^) and growth factors such as TGF-β from the degraded bone matrix ^7,24,25^, which further promotes tumor progression and osteolysis. Metastatic tumor cells can also acquire calcium from osteoblasts via gap junctions ^14^.

Additionally, glutamine produced by osteoclasts helps metastatic cancer cells overcome oxidative stress ^26^. Despite these progress in our understanding of metabolic adaptation during bone metastasis, further research is needed to understand how tumor cells obtain other essential nutrients from the bone microenvironment to support their survival and metastatic outgrowth.

In addition to acquiring stromal support, phenotypic plasticity enables certain tumor cells to adopt functions similar to those of stromal cells, enhancing their ability to adapt to foreign environments ^27^. For instance, glioblastoma cells can express endothelium-associated genes and form vessel-like structures, a process known as vasculogenic mimicry ^28–31^. Similarly, breast and prostate cancer cells can assume an osteoblast-like phenotype, producing bone matrix proteins and paracrine factors that disrupt the osteoblast–osteoclast balance, thereby manipulating the bone environment to their advantage ^32–36^. Precise characterization of tumor-stromal interactions is critical for identifying new forms of host-organ mimicry. Understanding these adaptive mechanisms may lead to the discovery of biomarkers predictive of organ-tropic metastasis and the development of novel therapeutic strategies.

In this study, we employed an *in vivo* metastatic niche labeling system ^37^ combined with single-cell RNA sequencing to systematically characterize the hematopoietic cells within the bone metastatic niche. Our analysis revealed a distinct population of macrophages enriched in the metastatic niche. These specialized macrophages typically maintain iron homeostasis and support red blood cell production in the erythroblastic islands (EBIs) of healthy bone marrow. We demonstrated that metastatic breast cancer cells hijack these iron-rich macrophages to facilitate their survival and growth within the bone.

Additionally, tumor cells develop a mechanism to mimic erythroblasts in response to hypoxic stress in the bone marrow. This hijacking of iron-rich macrophages by bone-metastatic tumors, at the expense of erythropoiesis, may contribute to metastatic cancer-associated anemia. Notably, similar iron metabolism features were observed in a subset of macrophages in human bone metastases across multiple cancer types. Our study provides a molecular mechanism linking the pathogenesis of bone metastasis with associated hematological complications, such as anemia.

## Results

### A unique macrophage population is enriched in the bone metastatic niche

To systematically characterize the cells spatially enriched in the bone metastatic microenvironment, we adopted an *in vivo* metastatic niche labeling system ^37,38^. Briefly, E0771, a mouse mammary tumor cell line derived from a spontaneous mammary gland tumor of a C57BL/6 mouse ^39,40^, was engineered to stably express secreted lipid permeable mCherry (sLP-mCherry) in addition to enhanced green fluorescent protein (EGFP) and firefly luciferase (**Figure 1A**). The lipid-permeable (LP) tag fused with mCherry enables only cells in close proximity to the tumor cells to capture the secreted mCherry from E0771 tumors ^37^. E0771 cells expressing sLP-mCherry, EGFP, and luciferase were injected into syngeneic C57BL/6 mice via the caudal artery to establish bone metastases (**Figure S1A**). Upon the development of bone metastasis, we observed that the majority of mCherry-labeled cells were within a maximum distance of 80 µm from the tumor cells, with the mCherry signal intensity decreasing with distance (**Figure 1B**-**C**, **S1B**), consistent with previous reports ^37^.

**Figure 1.**
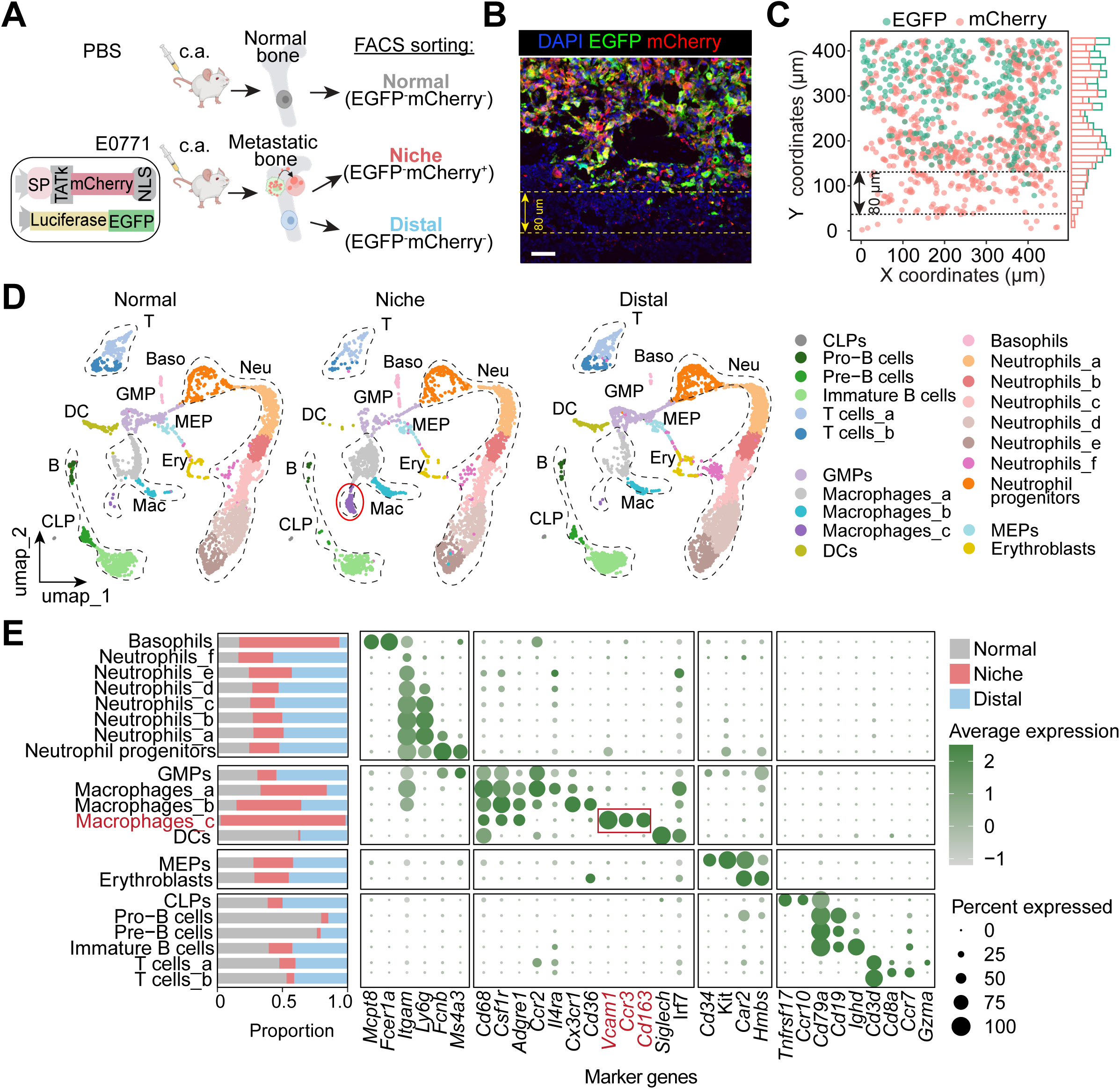
Discovery of a unique macrophage population in bone metastatic niche through niche labeling and scRNA-seq. (**A**) Schematic of niche labeling system and overview of the protocol. E0771 mouse mammary tumor cell line was engineered to stably express a firefly luciferase-IRES-EGFP reporter and secreted lipid permeable mCherry (sLP-mCherry) in which mCherry was fused to a signal peptide (SP) and a lipo-permeable Transactivator of Transcription (TATk) peptide. Double-labelled E0771 cells were injected into the caudal artery of 8- to 10-week-old female B6 albino mice to induce bone metastasis. Bone marrow cells were flushed from the femur and tibia and niche cells (EGFP^-^mCherry^+^) and distal cells (EGFP^-^mCherry^-^) were isolated by FACS. Normal bone marrow cells (EGFP^-^mCherry^-^) were also collected from healthy control mice sham-injected with PBS. (**B**) Representative immunofluorescent images of sLP-mCherry/EGFP labeled-E0771 cells within bone metastases, EGFP (green), mCherry (red); DAPI (blue), (**C**) Distribution of mCherry and EGFP signals from panel (B). (**D**) Uniform manifold approximation and projection (UMAP) plots showing unbiased clustering of cells isolated from normal (EGFP^-^mCherry^-^), niche (EGFP^-^mCherry^+^) or distal (EGFP^-^mCherry^-^) bone marrow following scRNA-seq integrated analysis. Red circle highlights the niche-specific cluster, Mac_c. (**E**) Bar plot showing the proportion of normal, niche and distal cells in each cluster. Bubble plot showing marker genes of each cluster. Dot size indicates the fraction of expressing cells, colored according to z-score normalized expression levels. CLP: Common Lymphoid Progenitor, T: T cell, B: B cell, DC: Dendritic Cell, GMP: Granulocyte-Macrophage Progenitor, MEP: Megakaryocyte-Erythroid Progenitor, Ery: Erythroblast, Mac: Macrophage, Baso: Basophil, Neu: Neutrophil. Scale bar, 50 μm in (B).

Using fluorescence-activated cell sorting (FACS), we isolated EGFP^-^mCherry^+^ niche cells and EGFP^-^mCherry^-^ distal cells from pooled metastatic bones (Figure S1C, right panel). As a control, we sorted an equivalent number of EGFP^-^mCherry^-^ normal bone marrow cells from pooled healthy bones (**Figure S1C**, left panel). To ensure better viability of hematopoietic cells, enzyme dissociation was not performed, thus mesenchymal, stromal, and endothelial cells were not included in this analysis. The niche, distal, and normal bone marrow cells were subsequently processed for single-cell RNA sequencing (scRNA-seq) to analyze their transcriptional profiles.

We annotated cell types in the scRNA-seq data using the Tabula Muris single-cell transcriptomic atlas of mouse bone marrow ^41^ as a reference. As expected, 21 distinct hematopoietic cell types were identified (**Figure 1D, S1D, and S1E**). We observed an imbalanced distribution of lymphoid and myeloid lineages between the niche, distal, and normal bone marrow. Specifically, there was a decreased proportion of T cells, B cells, and dendritic cells (DCs) in the metastatic bone marrow, particularly within the mCherry^+^ niche cells (**Figure 1E**), suggesting an immunologically “cold” phenotype. This observation aligns with clinical reports indicating poor responses to immune checkpoint inhibitors in patients with bone metastasis ^42^.

Conversely, the frequencies of granulocyte-lineage cells were increased in the metastatic bone marrow compared to normal bone marrow. Granulocyte-monocyte precursors (GMPs), neutrophil progenitors, and mature neutrophils were more prevalent in the distal metastatic bone environment, whereas basophils predominantly infiltrated the tumor niche (**Figure 1E**). Macrophages also showed increased presence in the metastatic bone marrow. Within this context, macrophage subsets exhibited distinct distributions: Mac_c and Mac_a were more enriched in the niche, while Mac_b was comparably distributed in both niche and distal areas (**Figure 1E, S1, S1F and S1G**).

All three macrophage subsets expressed macrophage markers *Cd68*, *Csf1r*, and *Adgre1* (encoding F4/80) (**Figure 1E**). Notably, in addition to macrophage markers, Mac_a exhibited high *Ccr2* and *Il4ra* expression, while Mac_b showed high *Cx3cr1* and *Cd36* expression. Mac_c specifically expressed *Vcam1*, *Cd163*, and *Ccr3* (**Figure 1E, S1H**).

Among all the cell types, the *Vcam1*^+^*Cd163*^+^*Ccr3*^+^ macrophage sub-cluster was found to be the most frequently enriched in the niche cells, but not in distal or normal bone marrow cells (**Figure 1D and 1E**). These prominent changes prompted us to further explore the properties and functions of this unique macrophage subpopulation.

### Niche-specific macrophages display heme and iron metabolism signature

Flow cytometry analysis revealed that VCAM1^+^CD163^+^CCR3^+^ cells are present in normal bone marrow, comprising about 1% of all live cells. The percentage of this population was elevated in the metastatic bone marrow of mice bearing bone metastasis of E0771 cells, and more significantly in bone lesions produced by Py8119, another mouse mammary tumor cell line derived from the MMTV-PyMT transgenic model of mammary gland tumors ^43,44^ (**Figure 2A**). Immunofluorescence imaging of the metastatic bone showed VCAM1, CD163, and CCR3-expressing cells localized near EGFP^+^ tumor cells (**Figure S1I and S1J**). This result, together with the finding of significant enrichment of the Mac_c population in the niche versus distal area (**Figure 1E and S1E**) suggests a dramatic shift of this population toward close proximity of metastatic tumors in addition to the modest increase of this population in tumor-bearing bone. This assessment not only validates that the sLP-mCherry labeling system effectively marks the neighboring cells, but also demonstrates that proximity labeling enables the discovery of rare cell populations enriched in specific niche areas within a diverse environment, a capability not feasible with scRNA-seq alone without specific labeling of niche cells.

**Figure 2.**
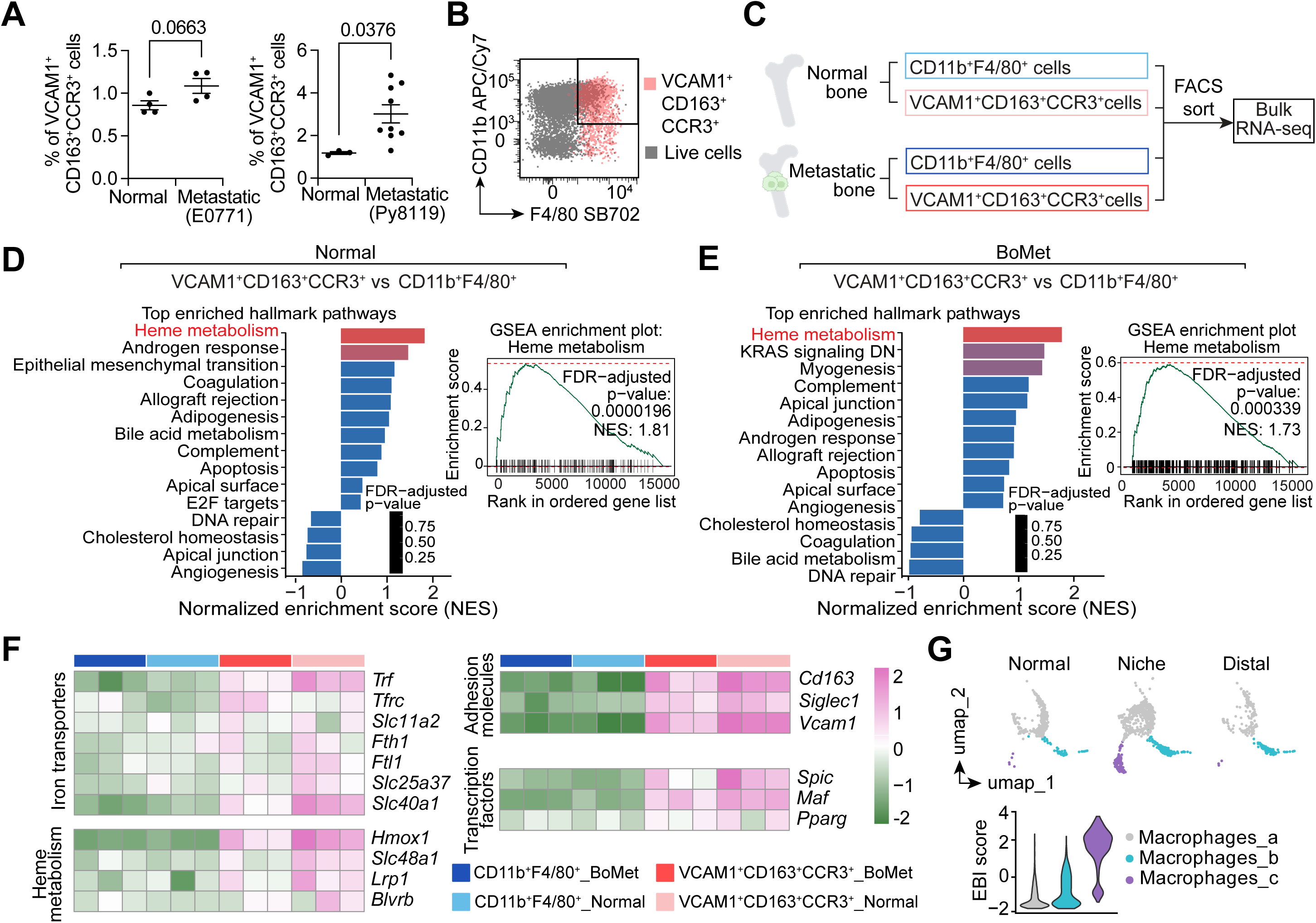
Identification of niche-specific macrophages as EBI macrophages. (**A**) Flow cytometry analysis of VCAM1^+^CD163^+^CCR3^+^ cell percentages in normal bones (n=4) and metastatic bones following c.a. injection of E0771 (n=4) or Py8119 (n=8) mouse mammary tumor cells into 8- to 10-week old female B6 albino mice. (**B**) Representative flow cytometry plots showing CD11b and F4/80 expression in VCAM1^+^CD163^+^CCR3^+^ cells versus total live bone marrow cells. (**C**) Schematic of bulk RNAseq workflow. VCAM1^+^CD163^+^CCR3^+^ or CD11b^+^F4/80^+^ cells were sorted from normal or metastatic bone following c.a. injection of E0771. n=3 biological replicates for each group. (**D** and **E**) GSEA showing the top differentially enriched hallmark pathways when comparing VCAM1^+^CD163^+^CCR3^+^ cells to CD11b^+^F4/80^+^ cells from normal bone marrow (D) and metastatic bone marrow (E), colored according to FDR-adjusted p-value. All differentially expressed genes were pre-ranked for GSEA using the hallmark gene set from MSigDB. P-values were adjusted using the Benjamini-Hochberg method. (**F**) Heatmap showing normalized expression of selected genes in VCAM1^+^CD163^+^CCR3^+^ and CD11b^+^F4/80^+^ cells from normal and metastatic bone marrow, colored according to z-score normalized expression levels. (**G**) UMAP showing three macrophage clusters in scRNA-seq, from niche, normal and distal bone marrow. Violin plot showing EBI score in the indicated macrophage clusters. The EBI score was defined by averaging expression of genes related to EBI functions, as listed in (F). EBI, erythroblastic island. GSEA, gene set enrichment analysis. All data is shown as mean ± SEM. Two-tailed unpaired Student’s t test was used to determine the statistical significance.

Next, we explored the identity of these niche-specific VCAM1^+^CD163^+^CCR3^+^ cells using flow cytometry. First, we confirmed that VCAM1^+^CD163^+^CCR3^+^ cells expressed myeloid lineage markers CD11b, F4/80, MHC2, and Ly6C, but not lymphoid markers CD79a, CD3 and CD8 (**Figure S1K and S1L**), consistent with our scRNA-seq annotation.

Furthermore, VCAM1^+^CD163^+^CCR3^+^ cells were verified as a sub-population of CD11b^+^ F4/80^+^ macrophages (Figure 2B). We further characterized the transcriptional landscape of the VCAM1^+^CD163^+^CCR3^+^ cells and CD11b^+^ F4/80^+^ macrophages from both normal and metastatic bone marrow using FACS followed by bulk RNA-seq (**Figure 2C, S2A, and S2B**). First, we confirmed that the top signature genes of the Mac_c cluster from the scRNA-seq, including *Vcam1*, *Cd163* and *Ccr3*, are highly expressed in the sorted VCAM1^+^CD163^+^CCR3^+^ cells, but not in CD11b^+^ F4/80^+^ macrophages (**Figure S2C**).

The heatmap of the top variable genes showed that samples primarily clustered based the cell types, but not the conditions (**Figure S2D**), suggesting the identity of the cells did not change under the metastatic condition.

To discover the unique features of the VCAM1^+^CD163^+^CCR3^+^ population, we compared their gene expression with that of CD11b^+^F4/80^+^ macrophages under the same conditions. Notably, gene set enrichment analysis (GSEA) of the differentially expressed genes revealed that, compared to CD11b^+^F4/80^+^ macrophages, VCAM1^+^CD163^+^CCR3^+^ cells are enriched for heme metabolism signature in both normal and metastatic conditions (**Figure 2D and 2E**). Heme metabolism signature was also enriched when we compared gene expression of Mac_c with Mac_a and Mac_b from our scRNA-seq data (**Figure S2E**).

There is a specialized population of resident macrophages in the bone marrow EBIs that serve as iron-rich nurse cells, supporting erythropoiesis ^45,46^. Previous pioneering work by multiple groups has characterized the key functional genes expressed by EBI macrophages ^47^. Specially, EBI macrophages supply iron for hemoglobin synthesis in red blood cells (RBCs) through the expression of iron transporters, which includes the ferric iron (Fe³⁺) transporter transferrin (encoded by *Trf*) and transferrin receptor (CD71 or TfR1, encoded by *Tfrc*), the non-heme ferrous iron (Fe²⁺) transporter DMT1 (encoded by *Slc11a2*), the iron storage proteins H-ferritin (encoded by *Fth1*) and L-ferritin (encoded by *Ftl1*), the mitochondrial iron transporter Mitoferrin-1 (encoded by *Slc25a37*), and the iron exporter ferroportin (FPN, encoded by *Slc40a1*). EBI macrophages also express genes involved in heme metabolism and iron recycling, including the heme degradation enzyme heme oxygenase-1 (HO-1, encoded by *Hmox1*), Biliverdin Reductase B (encoded by *Blvrb*), the heme transporter HRG1 (encoded by *Slc48a1*), and LRP1 (encoded by *Lrp1)*. These iron transporters and heme metabolism–associated genes showed significantly higher expression in the VCAM1^+^CD163^+^CCR3^+^ niche macrophage population compared to CD11b^+^F4/80^+^ macrophages (**Figure 2F**). *Cd163* and *Vcam1*, the two markers we identified, along with *Siglec1,* are key adhesion molecules critical for the interaction between EBI macrophages and erythroblasts ^45^. These molecules were significantly upregulated in the sorted VCAM1^+^CD163^+^CCR3^+^ cells, compared to CD11b^+^F4/80^+^ macrophages (**Figure 2F**). Additionally,VCAM1^+^CD163^+^CCR3^+^ cells exhibited high expression of key transcription factors involved in the development of EBI macrophages, including *Pparg* ^48^, *Maf* ^45^, and *Spic* ^49^ (**Figure 2F**). Here, we defined the EBI score by averaging expression of the genes listed above that are related to EBI functions. Surveying the EBI score across all macrophage clusters in our scRNA-seq data revealed that Mac_c showed the highest EBI score (**Figure 2G**). These findings suggest that VCAM1^+^CD163^+^CCR3^+^ macrophages, which are enriched in the bone metastatic niche, resemble EBI macrophages in recycling iron and maintaining iron homeostasis at the transcriptional level.

### Iron-rich macrophages are enriched at the tumor-stroma interfaces

FPN is the only known vertebrate iron exporter ^50^. Transferrin receptor CD71 is essential for iron uptake through binding transferrin and facilitating the endocytosis and release of iron into the cell ^46^. Immunofluorescence imaging of metastatic bone sections revealed that FPN was primarily expressed in F4/80^+^ macrophages (**Figure 3A**).

**Figure 3.**
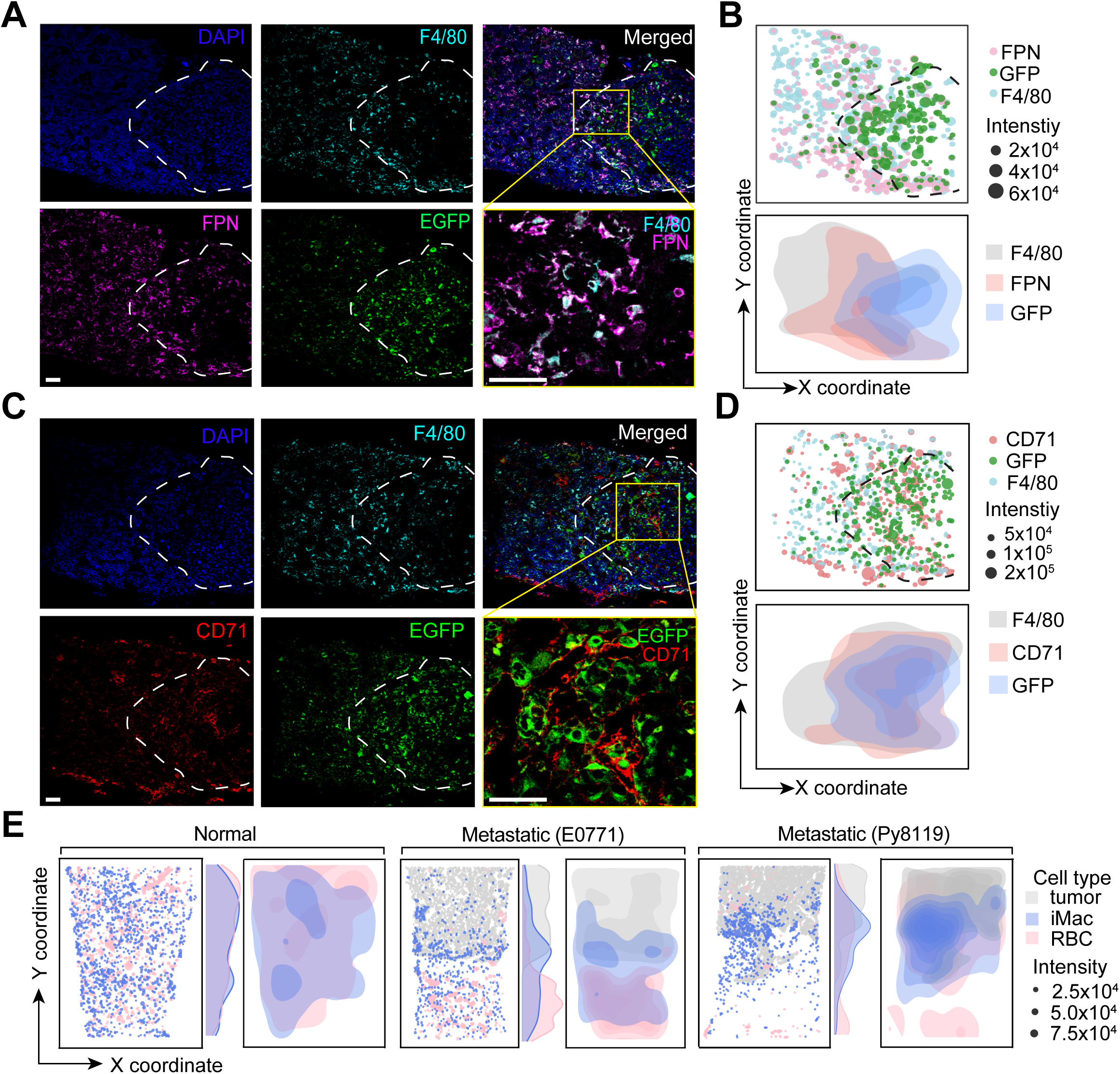
Iron-rich macrophages are enriched at the tumor-stroma interface. (**A**) Representative immunofluorescent images of F4/80 (macrophage marker, cyan), FPN (iron exporter, magenta), and EGFP (tumor cell, green) in E0771 bone metastases. The white dashed lines indicated tumor-stroma edge. (**B**) Scatter plot (top panel) and density plot (bottom panel) showing iron exporter (FPN^+^) expressing macrophages (F4/80^+^) enriched at the edge of tumor (EGFP^+^). The black dashed lines indicated tumor-stroma edge. (**C**) Representative immunofluorescent images of F4/80 (macrophage marker, cyan), CD71 (iron uptake gene, red), and EGFP (tumor cell, green) in E0771 bone metastases. The white dashed lines indicated tumor-stroma edge. (**D**) Scatter plot (top panel) and density plot (bottom panel) showing tumor cells (EGFP^+^) expressed iron uptake gene CD71. The black dashed lines indicated tumor-stroma edge. (**E**) Scatter plot and density plot image of Prussian blue-stained iron-rich macrophages (iMac) and red blood cells (RBCs) in normal bone and bone metastases with E0771 or Py8119 cells. Marginal density plots showing the distribution of iMac, RBCs and tumor cells along y-coordinate. See Supplementary Figure S3, A to C for raw and split-channel images. Scale bars in (A) and (C), 50 µm.

Quantification of images suggested an increased presence of FPN^+^F4/80^+^ macrophages at the tumor-stroma interfaces (**Figure 3A and 3B**). Meanwhile, the iron uptake gene CD71 was predominantly expressed in EGFP^+^ tumor cells (**Figure 3C and 3D**). These findings suggest a potential iron flow from macrophages to tumor cells within the bone metastatic environment.

To further examine the distribution of iron-rich macrophages in bone marrow, we performed Prussian blue staining ^51,52^ on normal and metastatic bone sections. In this assay, only ferric iron stored as ferritin and hemosiderin stains bright blue, while iron bound to porphyrin (forming heme) or free labile iron (Fe^2+^) inside cells cannot be detected. RBCs appear brown in the stained sections ^51^. Tumor boundaries can be recognized based on cell morphology and the EGFP signal of tumor cells from adjacent slides. Image quantification showed an even distribution of iron-rich macrophages and RBCs in normal bone marrow (**Figure 3E and S3A**). However, in metastatic bone, we observed a significant enrichment of iron-laden macrophages at the edge of E0771 tumor (**Figure 3E and S3B**), consistent with our observations of VCAM1^+^CD163^+^CCR3^+^ cells (**Figure S1I and S1J**) and FPN^+^F4/80^+^ macrophages (**Figure 3A**). Analysis of Py8119 metastatic bone showed a similar phenomenon (**Figure 3E and S3C**) to what was observed in E0771. Taken together, our findings indicate that tumors disrupt the normal distribution of EBI macrophages in the bone marrow, leading to the enrichment of iron-rich macrophages in tumor-adjacent areas, away from their normal enrichment in EBIs. This redistribution may impair the macrophages’ role in supporting erythropoiesis.

### Red blood cell formation is impaired in metastatic bone marrow

Next, we investigated whether erythropoiesis was impaired in bone marrow with metastatic tumors. Mature red blood cells (erythrocytes) are produced from progenitor cells through four nucleated erythroblast stages: proerythroblast (Pro-E, stage I), basophilic erythroblast (Baso-E, stage II), polychromatic erythroblast (Poly-E, stage III), and orthochromatic erythroblast (Ortho-E, stage IV). This is followed by the formation of immature red blood cells (reticulocytes, stage V) and mature red blood cells (erythrocytes, stage VI) ^53,54^ (**Figure 4A**). We quantified cells at each stage of erythroid differentiation in normal and metastatic murine bone marrow based on the expression levels of TER119, CD44, and cell size using flow cytometry ^53^ (**Figure 4B**). We observed a decreased count of reticulocytes (V) and erythrocytes (VI) in the metastatic bone marrow (**Figure 4C**), consistent with our observation in the Prussian blue stained sections (**Figure 3E**). Although the total count of nucleated erythroblasts (I+II+III+IV) did not change (**Figure 4C**), the ratio of Pro:Baso:Poly:Ortho was 1:1.1:5.6:14 in metastatic bone marrow, compared to a ratio of 1:2.5:4.4:7.5 in normal bone marrow, which typically follows a 1:2:4:8 pattern ^53^ (**Figure 4D**). This suggests disrupted erythropoiesis in the metastatic bone marrow. When extracellular iron levels are low, cells increase the expression of CD71 to enhance iron uptake, mediated by the iron-regulatory proteins (IRPs) ^55,56^. Significantly increased CD71 levels were indeed observed across different stages of erythroid cells in metastatic bone marrow (**Figure 4E and 4F**), indicating insufficient iron availability in the local environment. Similar observations were noted in Py8119 metastatic bone marrow (**Figure S4A to S4E**).

**Figure 4.**
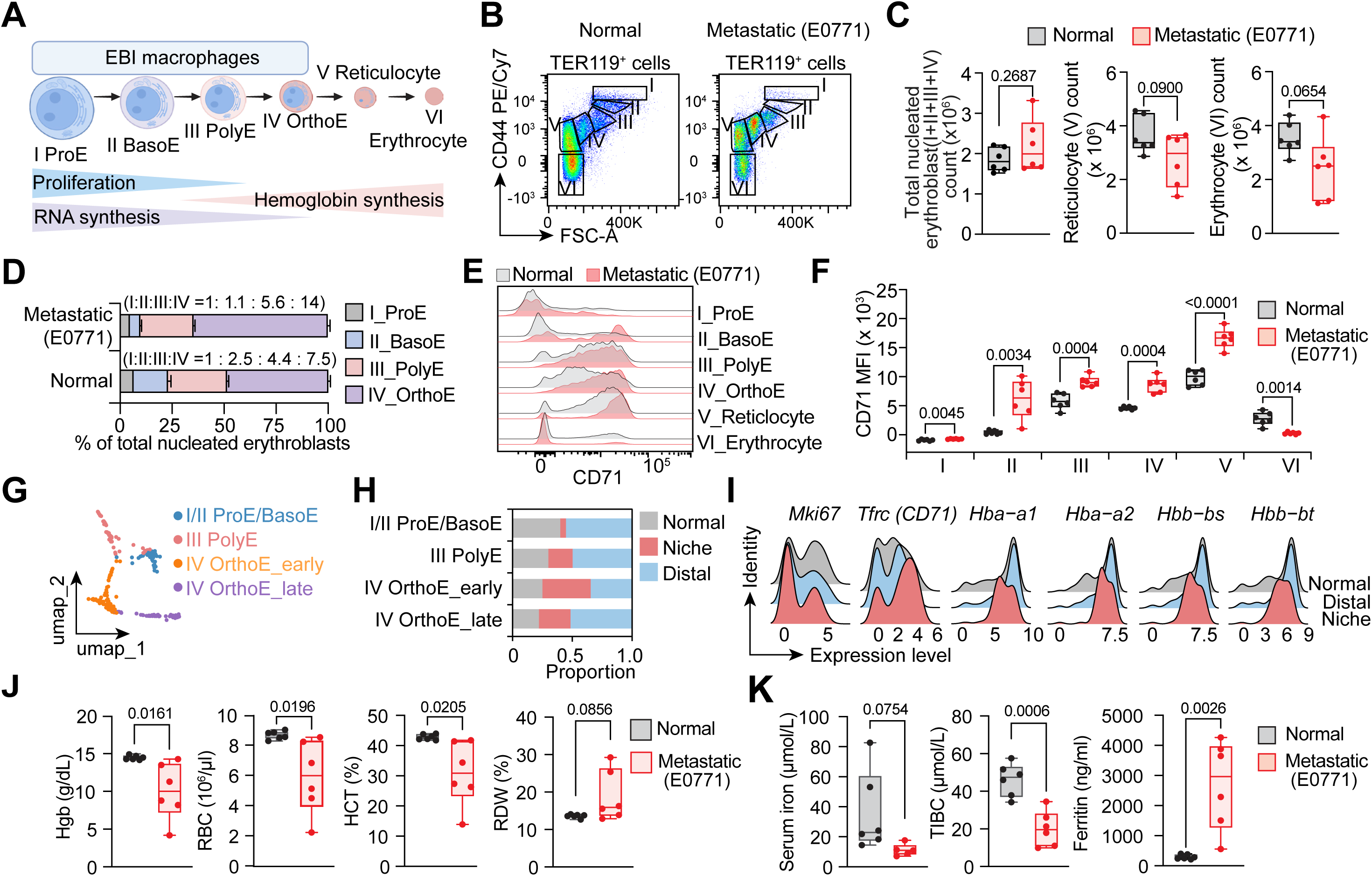
Inefficient erythropoiesis in the metastatic bone marrow leads to anemia. (**A**) Schematic of erythroid lineage cell differentiation. (**B**) Representative flow cytometry plots showing different stage of erythroid cells based on CD44 expression and cell size in CD45^-^TER119^+^ cells from normal and E0771 metastatic bone marrow. (**C**) Quantification of the proportion of total nucleated erythroblast stages (I-IV), reticulocyte (V) and erythrocyte (VI) in normal or E0771 metastatic bone marrow. n=6 biological replicates for each group. (**D**) Bar plot showing the proportions of ProE (I), BasoE (II), PolyE (III), and OrthoE (IV) within the total nucleated erythroblast population. (**E** and **F**) Representative flow cytometry histograms (E) and quantification (F) of CD71 expression in different stage of erythroid cells. n=6 biological replicates for each group. (**G**) UMAP plot of unbiased sub-clustering of erythroblasts. (**H**) Bar plots showing the proportion of each erythroblast sub-cluster in normal, niche and distal bone marrow. (**I**) Ridge plots showing expression distributions of selected genes in erythroblasts from normal, niche and distal bone marrow. (**J** and **K**) Quantification of Hemoglobin (Hgb) level, RBC count, Hematocrit (HCT), and RBC distribution width (RDW) in the blood (J) and iron level, total iron binding capacity (TIBC) and ferritin level in the serum (K) from E0771 bone metastasis mice or normal control mice. n=6 biological replicates for each group. All data is shown as mean ± SEM. Two-tailed unpaired Student’s t test (C, J, K), and multiple unpaired Student’s t test (F) were used to determine the statistical significance.

Next, we leveraged our scRNA-seq data to explore the number and gene expression of different stages of erythroblasts in the metastatic bone niche. We annotated the sub- clustered erythroblasts based on a scRNA-seq transcriptome analysis of erythroblasts from human bone marrow ^57^ (**Figure 4G, S4F, and S4G**). Consistent with our flow cytometry results, we noted a significant decrease in early-stage erythroblasts (I+II+III) in the metastatic niche cells compared to distal and normal bone marrow cells (**Figure 4H and S4F**), which suggest erythroblasts may fail to compete with tumor cells for macrophage support within the tumor niche. We observed increased *Trfc* (CD71) expression and decreased expression of globin genes, *Hba-a1*, *Hba-a2*, *Hbb-bs*, and *Hbb-bt*, in erythroblasts from the tumor niche, while the proliferation marker Mki67 remained unchanged, compared to those from normal or distal bone marrow (**Figure 4I**). This data further confirms a defect in erythroid differentiation and a deprivation of iron availability to the erythroblasts within the tumor niche.

We observed decreased hemoglobin (Hgb) levels, reduced RBC counts, decreased RBC per unit volume (measured as Hematocrit, HCT), and increased variation in RBC size (measured as red blood cell distribution width, RDW) in the blood of mice with bone metastasis (**Figure 4J**). Additionally, we noted decreased circulating serum iron concentration and total iron-binding capacity (TIBC), despite elevated serum ferritin levels (**Figure 4K**). These findings are consistent with clinical manifestation of cancer-associated anemia ^58,59^.

Together, our data suggest that erythropoiesis supported by EBI macrophages was impaired in metastatic bone, contributing to a reduced Hgb level and red blood cell count in the blood, as well as other general features of anemia.

### Depletion of *Spic*-expressing iron-recycling macrophages impedes bone colonization

We next investigated the role of this macrophage subpopulation in regulating metastatic tumor growth. Previous research indicated that the transcription factor *Spic* is essential for the development of EBI macrophages in the bone marrow ^49^. First, we found that Mac_c specifically showed high transcriptional expression of *Spic,* along with high expression of its downstream target gene *Slc40a1* (FPN) ^60^ (**Figure 5A**). Using Spic- EGFP reporter mice, which allow tracking of endogenous Spic protein expression, we confirmed via flow cytometry and immunofluorescence that Spic-EGFP^+^ cells are VCAM1^+^CD163^+^CCR3^+^ (**Figure 5B, 5C and S5A**). Furthermore, Spic-EGFP^+^ cells were enriched at the tumor-stroma interfaces in metastatic bone (**Figure S5B**). Importantly, co-culture of Spic-EGFP^+^ macrophages with tumor cells promoted tumor growth *in vitro*, compared to co-culturing with Spic-EGFP^-^ macrophages (**Figure 5D and 5E**).

**Figure 5.**
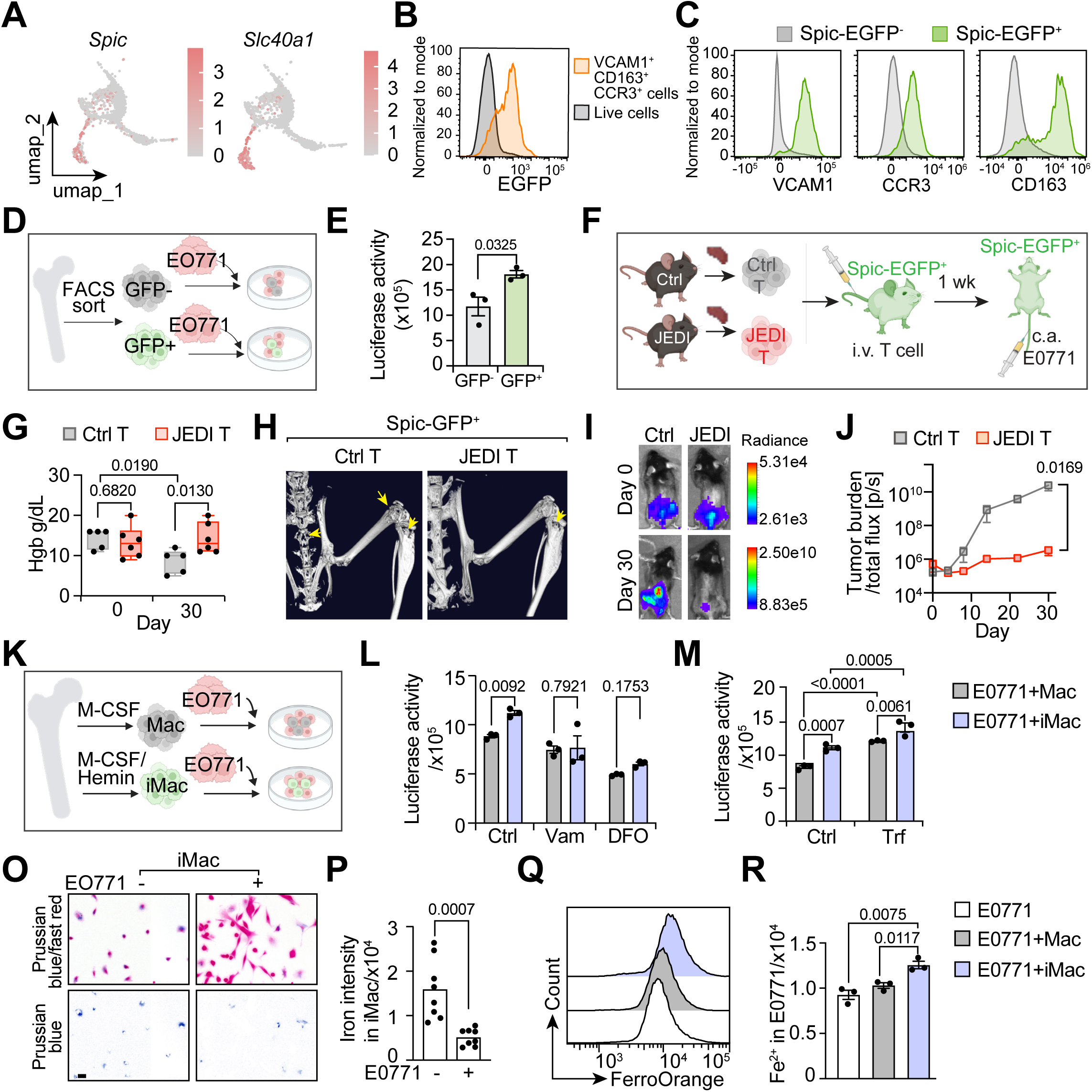
Iron-rich macrophage support tumor growth. (**A**) UMAP plots showing *Spic* and *Slc40a1* (FPN) expression in macrophage clusters, colored according to z-score normalized expression levels. (**B**) Representative flow cytometry histograms showing expression of Spic-EGFP reporter in VCAM1^+^CD163^+^CCR3^+^ cells or bone marrow live cells. (**C**) Representative flow cytometry histograms showing expression of VCAM1, CCR3, and CD163 in Spic-EGFP^-^ or Spic-EGFP^+^ cells. (**D**) Schematic of E0771 co-culture with Spic-EGFP^-^ or Spic-EGFP^+^ macrophages. (**E**) Luciferase activity representing tumor cell growth when co-cultured with sorted Spic-EGFP^-^ or Spic-EGFP^+^ macrophages. n=3 biological replicates. (**F**) Schematic of the delivery of JEDI T cells or control T cells via intravenous injection (i.v.) one week prior to E0771 tumor cell injection through caudal artery (c.a.) in 8- to 12- week-old female Spic-EGFP^+^ mice. (**G**) Hemoglobin (Hgb) level from day 0 (1 week after T cell injection) and day 30 post-tumor inoculation, in Spic-EGFP^+^ mice injected with Ctrl or JEDI T cells. n=5 biological replicates. (**H**) Representative microCT scans of bones from Spic-EGFP^+^ mice injected with Ctrl or JEDI T cells, collected on day 30 post-tumor inoculation. Bone metastases indicated by yellow arrow. (**I**) Representative bioluminescence imaging (BLI) images of Spic-EGFP^+^ mice injected with Ctrl or JEDI T cells, on day 0 and day 30 post-tumor inoculation. (**J**) Bone metastasis burden quantified by BLI (I) from day 0 to day 30 post-tumor inoculation, in Spic-EGFP^+^ mice injected with Ctrl or JEDI T cells. n=6 replicates for JEDI T group and n=6 replicates for Ctrl T group. (**K**) Schematic of E0771 co-culture with M-CSF-induced macrophage (Mac) or M-CSF/Heme-induced iron-rich macrophages (iMac). (**L** and **M**) Luciferase activity showing tumor growth when co-cultured with Mac or iMac, treated with iron chelator Deferoxamine (DFO) and the iron exporter FPN inhibitor vamifeport (Vam) (L) or treated with iron-binding protein transferrin (M). n=3 replicates for each group. (**O**) Reprehensive bright-field images of iMac alone or iMac co-cultured with E0771. Top: Prussian blue stain with fast red nuclear stain, bottom: Prussian blue stain only. (**P**) Quantification of iron stored in macrophage detected by Prussian blue staining. n=8 replicates. (**Q**) Representative flow cytometry histograms of FerroOrange staining in E0771 cells when cultured alone, with Mac or with iMac. (**R**) Quantification of free iron level in tumor cell detected by mean fluorescence intensity (MFI) of FerroOrange in E0771 cells. n=3 replicates for each group. Scale bar, 10 μm in (**O**). All data is shown as mean ± SEM. Two-tailed unpaired Student’s t test (E, P, R), and Two-Way ANOVA (I, J, L, M) were used to determine the statistical significance.

To deplete the Spic-EGFP^+^ population *in vivo*, we utilized the genetically modified JEDI (Just EGFP Death-Inducing) mouse model ^61^. In JEDI mice, T-cells are engineered to express a TCR that recognizes EGFP, enabling selective depletion of EGFP-expressing cells upon transfer of JEDI T-cells into EGFP reporter animals ^61^. We injected equal numbers of control CD8^+^ T cells or JEDI CD8^+^ T cells into Spic-EGFP^+^ mice one week before tumor injection (**Figure 5F**). Depletion of Spic-EGFP^+^ cells for one week did not affect Hgb levels (**Figure 5G**). Subsequently, mCherry^+^ E0771 tumor cells were administered via either caudal artery (**Figure 5F to 5J**) or intracardiac injection (**Figure S5C to S5F**) to generate bone metastasis. Depletion of Spic-EGFP^+^ cells slowed tumor growth and reversed the decline of hemoglobin levels observed 3-4 weeks after tumor implantation in the control group (**Figure 5G to 5J, S5D to S5F**). To confirm that the tumor-inhibitory effect was not caused by any off-target effect of JEDI T cells, we injected JEDI T cells into Spic-EGFP^+^ or Spic-EGFP^-^ littermates and found that tumor growth was impaired only in Spic-EGFP^+^ mice (**Figure S5G to S5I**), indicating that iron-recycling macrophages support tumor growth in the bone.

### Iron-rich macrophages support tumor growth through iron transportation

It is well known that iron is essential for tumor growth ^62^ , as it plays a critical role in the proliferation, cell-cycle control, and genome instability of cancer cells, such as maintaining the catalytic function of ribonucleotide reductase and DNA polymerases, and influencing the activity of the p53 tumor suppressor ^63,64^. The iron metabolism characteristics of this niche-specific macrophage population led us to hypothesize that these macrophages support tumor growth through iron transport. To test this hypothesis, we first induced bone marrow cells to differentiate into iron-rich macrophages (hereafter referred to as iMac) with heme (Hemin, Sigma) and M-CSF *in vitro* for 5 days (**Figure 5K**), as previously described ^49^. Macrophages (hereafter referred to as Mac) derived from bone marrow cells induced with M-CSF alone were used as a control. As expected, heme induced Spic-EGFP expression in macrophages (**Figure S5K**). iMac displayed a brown color after heme uptake (**Figure S5L**) and showed significantly higher iron storage, as indicated by Prussian blue staining, compared to M-CSF-induced macrophages (**Figure S5M**). Notably, when equal numbers of iMac and Mac were co-cultured with E0771 tumor cells, increased tumor growth was observed when tumor cells were co-cultured with iMac as compared to Mac (**Figure 5L and 5M**).

To determine if this tumor-promoting function is mediated by iron transportation, we blocked iron transport using the iron chelator Deferoxamine (DFO) and the iron exporter FPN inhibitor vamifeport (Vam). This intervention mitigated the tumor-promoting effects of iMac co-culture (**Figure 5L**). Furthermore, this tumor-promoting effect was enhanced by the addition of transferrin, which facilitates iron transport (**Figure 5M**). Similar observations were made when co-culturing iMac with 4T1 mouse mammary tumor cells (**Figure S5O to S5Q**). Prussian blue staining showed that ferric iron level in heme-induced iMac was decreased when iMac were co-cultured with tumors (**Figure 5O and 5P**). Free labile iron levels increased in tumor cells after co-culture with iMac, as evidenced by staining with the iron probe FerroOrange ^65^ (**Figure 5Q and 5R**). Taken together, our study suggests that iMac promotes tumor growth by supplying iron to the tumor cells.

### Bone metastatic tumor cells up-regulate heme synthesis and globin genes in response to hypoxia

We further investigated how iron transport by iMac influences tumor cell metabolism and survival within the bone metastatic microenvironment. To this end, we dissociated cells from both E0771-developed bone metastatic tumor and primary tumor for scRNA-seq (**Figure 6A and S6A**). GSEA revealed an upregulation of heme metabolism pathway in the metastatic tumor cells compared to tumor cells from the primary sites (**Figure 6B**). Heme synthesis and degradation involve the conjugation to or removal of iron from protoporphyrin through a series of enzymatic reactions ^66,67^. Our findings showed that multiple enzymes involved in heme synthesis are upregulated in tumors that metastasize to the bone, whereas *Hmox1*, which encodes for the enzyme that catalyzes the rate-limiting step in heme degradation, was down-regulated (**Figure 6C**). It is plausible that the presence of iron-rich macrophages in the bone niche provides the necessary iron source for bone metastatic tumors to synthesize heme.

**Figure 6.**
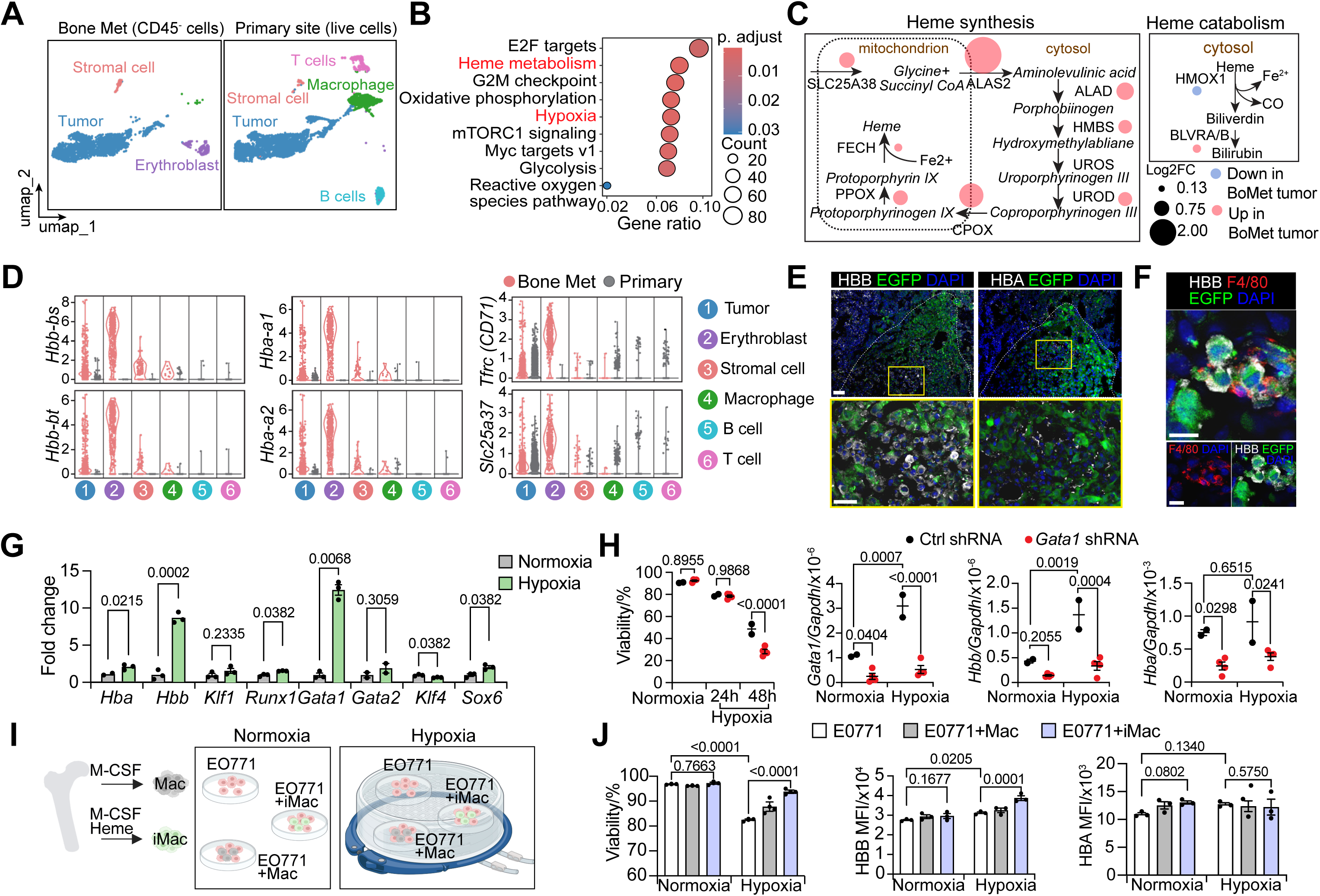
Up-regulated heme synthesis and globin gene expression in bone metastatic tumor in response to hypoxia. (**A**) UMAP plots showing unbiased clustering of CD45^-^ cells isolated from E0771 bone metastases (BoMet) and total live cells isolated from E0771 primary tumors after scRNA-seq analysis. (**B**) GSEA analysis showing differentially enriched hallmark pathways when comparing tumor cells from bone metastases with primary tumors in scRNA-seq. Dot represents the gene counts of enriched terms, colored according to adjusted p-value (p. adjust). The top 10 enriched pathways were identified using a p.adjust cutoff of 0.05 with the Benjamini-Hochberg method. (**C**) Bubble plots of heme- metabolism gene expression in E0771 cells from bone metastases compared with primary tumors in scRNA-seq. Dot size indicates log2 fold change, colored according to up- or down-regulation in bone metastases. (**D**) Violin plots showing expression of globin genes, iron uptake or utilization genes in different cell clusters from bone metastases or primary tumors. (**E**) Representative immunofluorescent images of HBA (white) or HBB (white), and EGFP (green) in EGFP^+^ E0771 bone metastases. Dashed lines indicated tumor-stroma edge. (**F**) Representative immunofluorescent images of HBB (white), F4/80 (red), and EGFP (green) in EGFP^+^ E0771 bone metastases. (**G**) Expression of globin genes and key transcription factors in E07771 when cultured under normoxia (20% O2) and hypoxia (2% O2) by RT–qPCR. Expression levels were normalized to normoxia condition. (**H**) Viability of 2 control and 4 *Gata1* shRNA knockdown E0771 cell lines cultured under normoxic and hypoxic conditions for 24 or 48 hours. Expression levels of the indicated genes were detected by RT–qPCR after 24 hours of hypoxic culture. See Supplementary Figure S6D for viability and gene expression data for the individual shRNAs. (**I**) Schematic of E0771 cells cultured alone, with Mac or with iMac, under normal or hypoxic condition. (**J**) Viability and expression of HBB and HBA in tumor cells cultured alone, with Mac or with iMac, under normal or hypoxic condition for 24h as measured by flow cytometry. n=3 replicates for each group. Scale bars, 50 μm in (E) and 10 μm in (F). All data is shown as mean ± SEM. Multiple unpaired Student’s t test (G), and Two-Way ANOVA (H, J) were used to determine the statistical significance.

Hemoglobin, composed of four globin subunits each with a heme, was previously considered to be expressed only by cells of the erythroid lineage ^68^. However, it has been shown to be present in multiple nonerythroid cells ^68^, including chondrocytes ^69^, neurons ^70^, alveolar epithelial cells ^71^, as well as circulating tumor cells (CTCs) from breast cancer patients ^72^ and 4T1 mouse breast cancer cells ^73^. Notably, in our scRNA-seq data, the iron uptake gene *Tfrc* (CD71) and the iron utilization gene *Slc25a37* (Mitoferrin-1) were expressed at similar levels in both metastatic and primary tumor cells. In contrast, the globin genes, *Hba-a1*, *Hba-a2*, *Hbb-bs* and *Hbb-bt*, were significantly expressed in E0771 cells in bone metastasis (**Figure 6D and S6B**). These findings suggest that iron metabolism is a common feature in tumor, whereas the biogenesis globin becomes more pronounced when tumors metastasize to the bone. At the protein level, we observed high expression of hemoglobin subunit beta (HBB, beta-globin), but not hemoglobin subunit alpha (HBA, alpha-globin) in the bone metastatic tumor cells by immunofluorescent staining (**Figure 6E**), suggesting a different regulatory mechanism between HBA and HBB at the mRNA and protein levels within the tumor population. HBB-expressing tumor cells were found to be in close contact with macrophages (**Figure 6F**), supporting the notion of macrophages playing a nursing role in the tumor microenvironment.

The bone is considered one of the most hypoxic organs in the body ^21^. As expected, our scRNA-seq analysis revealed upregulated hypoxia-related features in the bone metastasis tumor cells, compared to primary tumors (**Figure 6B**). Hypoxia is known to induce hemoglobin expression ^69,71^. Immunofluorescent staining confirmed the co-expression of HBB and HIF1A, the hypoxia-inducible transcription factor, in the bone metastatic E0771 cells *in vivo* (**Figure S6C**). We also observed the mRNA level of *Hbb was* significantly upregulated in the cultured E0771 cells under hypoxic environments, while *Hba* expression only slightly increased. To explore how globin gene expression in tumor cells was regulated, we examined the expression of *Klf1*, *Runx1*, *Gata1*, *Gata2*, *Klf4*, and *Sox6*, key transcription factors crucial for erythropoiesis, in E0771 cells under normoxic and hypoxic conditions (**Figure 6G**). We found that hypoxia induced a significant increase of *Gata1* expression in E0771 cells (**Figure 6G**). Short interfering RNA (shRNA)-mediated knockdown of *Gata1* led to reduced cell viability under hypoxic stress, as well as reduced *Hbb* and *Hba* mRNA expression (**Figure 6H and S6D**).

These results suggested GATA1 as a potential major regulator induced by hypoxia, controlling globin production in E0771 cells.

We next investigated whether iMac could contribute to globin gene expression under hypoxia. The iMac-tumor co-culture experiments were performed under normoxia or hypoxia condition (**Figure 6I**). First, we found hypoxia alone could induce HBB protein expression in E0771 cells, as demonstrated by intracellular staining via flow cytometry (**Figure 6J**, right panel). HBB expression was significantly increased in E0771 cells when co-cultured with iMac under hypoxia (**Figure 6J**, right panel). Cell viability assay also demonstrated that iMac confers protection to tumor cells against hypoxia-induced cell death, which was accompanied by elevated HBB expression (**Figure 6J**, left panel). This finding aligns with previous reports that the depletion of HBB in breast cancer cells increases cell apoptosis ^72^. Taken together, our data suggests macrophages supply iron to tumors, facilitating tumor growth and protecting them from hypoxia-induced cell death through the increased production of hemoglobin in tumor cells.

### Macrophages showed iron metabolism features and tumor cells expressed globin genes in human bone metastases

We explored the iron/heme metabolism features in scRNA-seq data of human bone metastases from a variety of primary cancers including breast cancer (n=9), lung cancer (n=6), kidney cancer (n=15), thyroid cancer (n=2), colorectal cancer (n=1), and esophageal cancer (n=1) along with healthy controls (n=5) (**Figure S7A**) ^74^. We first investigated whether the Mac_c population we discovered in the mouse bone metastatic environment also existed in humans. Notably, we observed the highest Mac_c signature score in one of the clusters in the human scRNA-seq data, which was identified as monocyte/macrophage in the source study ^74^ (**Figure S7B**). Most of the Mac_c signature genes showed specificity in the macrophage sub-population in both mouse (**Figure S7C**) and human (**Figure S7D**). Importantly, we found that Mac_c signature genes specifically expressed in a subset of *CD68*^+^ macrophage (**Figure 7A and 7B**) across multiple cancer types (**Figure 7B**), suggesting a widespread role of the iron metabolism features in the tumor microenvironment. *SLC40A1*, which encodes the only known iron exporter FPN, and *HMOX1*, which is critical for EBI macrophages to recycle iron, showed the highest expression in the same macrophage cluster (**Figure S7E**), suggesting that this macrophage subtype acts as the major iron supplier in the environment. We further confirmed the presence of macrophages with iron metabolism features in bone metastases by performing immunofluorescent staining for CD68 and FPN, along with Prussian blue staining, on bone metastasis sections from breast, kidney, and lung cancer patients (**Figure 7C**).

**Figure 7.**
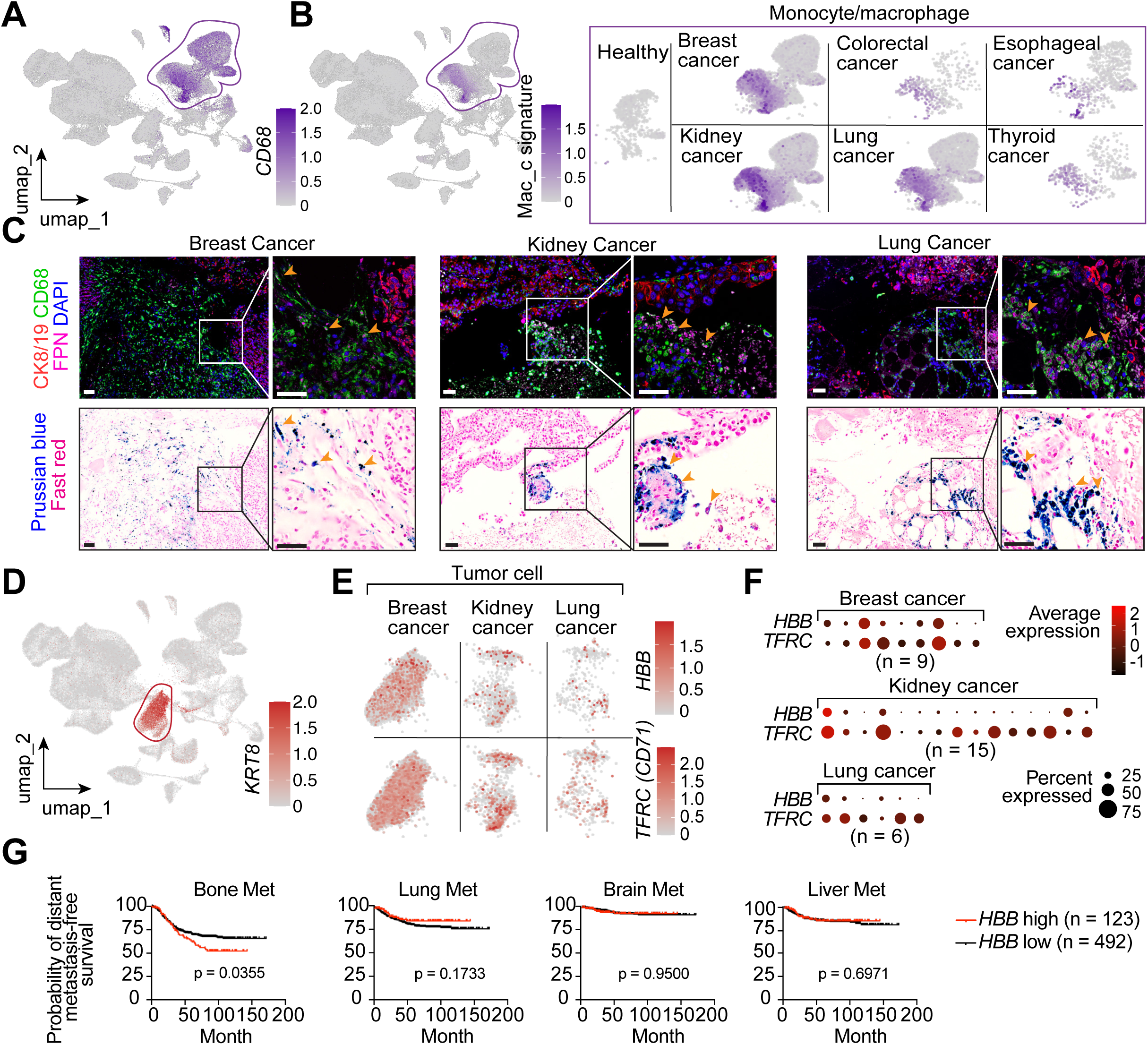
Validation of iron-rich macrophage in clinical samples. (**A** and **B**) UMAP plot showing expression of *CD68* (A) and Mac_c signature score (B) among 160,000 single cells from human bone metastases across six cancer types. See Supplementary Figure S7, A and B for cancer types and cell type annotation. Right panel in (**B**) showing Mac_c signature in monocytes/macrophages from different healthy control or cancer types. Colored according to z-score normalized expression levels. See Supplementary Figure S7, C and D for individual gene expression. (**C**) Immunofluorescent staining of tumor markers CK8/CK19, macrophage marker CD68, and iron exporter FPN, along with Prussian blue staining, in consecutive bone metastasis sections from breast, kidney, and lung cancer patients. Orange arrows indicate FPN^+^CD68^+^ cells. (**D** and **E**) UMAP plot showing expression of tumor/epithelium marker KRT8 among all cell types, and *HBB* and *TFRC* (CD71) in tumor cells from breast cancer, kidney cancer, and lung cancer (E). Colored according to z-score normalized expression levels. (**F**) Dot plot showing expression of *HBB* and *TFRC* (CD71) in tumor cells from individual breast cancer, kidney cancer, and lung cancer patients. Dot size indicates the percentage of cells expressing the gene, and color indicates the average expression level. (**G**) Kaplan-Meier plot showing the probability of cumulative bone, lung, liver, and brain metastasis-free survival based on *HBB* expression levels. The *HBB* high group (n=123) represents the top 20% of ranked expression, while the *HBB* low group (n=492) includes the remaining cases. Scale bars, 50 μm in (C).

Next, we examined the gene expression in the tumor cluster annotated by the source study ^74^, which was indicated by the expression of *KRT8* (**Figure 7D**). Of note, there were only sufficient cancer cells in breast cancer, lung cancer and kidney cancer patient samples for analysis (**Figure S7F**). *HBB* expression was detected in all three types of cancer cells (**Figure 7E**). More interestingly, we found a positive correlation between *HBB* and *TFRC* (CD71) expression (**Figure 7E and 7F**), consistent with the mouse data (**Figure S6B**). This observation indicates that iron uptake is coupled with beta-globin gene expression in cancer cells in human.

We next asked whether HBB expression correlated with bone metastasis. By analyzing data from the Erasmus Medical Centre/Memorial Sloan-Kettering Cancer Center (EMC- MSK) cohort ^10^, which includes 615 breast cancer patients, we found that high HBB expression in breast cancer was associated with higher risk of bone metastasis but not with lung, liver or brain metastasis (**Figure 7G**). Taken together, the presence of these iron-rich macrophages in human bone metastases, coupled with the expression of HBB in tumor cells and its association with bone metastasis, underscores the broad clinical relevance of our findings.

## Discussion

In this study, we discover the enrichment and functional role of *Vcam1*^+^*Cd163*^+^*Ccr3*^+^ EBI macrophages in the bone metastatic environment and reveal how the interactions with these macrophages stimulate erythroblast mimicry in tumor cells, thereby allowing them to adapt and thrive in the hypoxic bone marrow environment. Our study also elucidates a mechanism for cancer-associated anemia, offering critical insights for developing therapeutic strategies for both bone metastasis and cancer-associated anemia (**Figure 8**).

**Figure 8.**
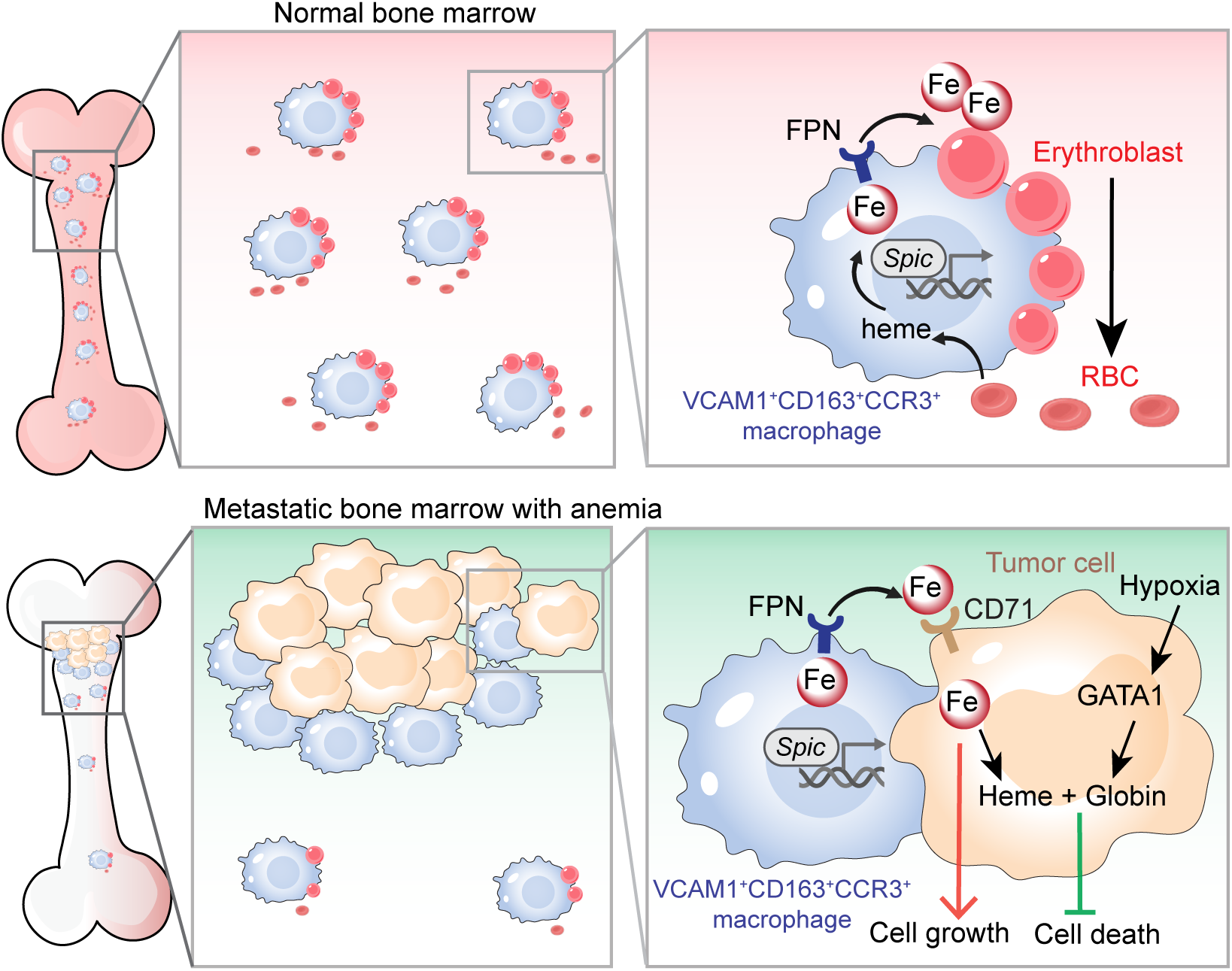
Tumor cells hijack EBI macrophages and mimic erythroblast to grow in bone. Model illustrating the interplay between breast cancer cells, EBI macrophages, and erythroblasts in the bone marrow. In normal bone marrow (upper panel), *Spic*- expressing VCAM1^+^CD163^+^CCR3^+^ EBI macrophages provide recycled iron from heme to erythroblasts via FPN, supporting hemoglobin synthesis and red blood cell production. In metastatic bone marrow (lower panel), tumor cells attract iron-rich VCAM1^+^CD163^+^CCR3^+^ macrophages away from EBIs, and import the recycled iron to support their growth and heme synthesis. In the meantime, tumor cells upregulate globin gene expression under the increased expression of GATA1 in response to hypoxia to better survive in the hypoxic bone environment. This leads to reduced availability of iron for erythroblasts, impairing erythropoiesis and contributing to the development of anemia.

Combining the *in vivo* niche labeling system with the high-resolution capabilities of scRNA-seq, we comprehensively profiled the diverse cellular populations in the complex bone marrow microenvironment. Such analysis led to the identification of *a Vcam1*^+^*Cd163*^+^*Ccr3*^+^ macrophage sub-population that is highly enriched in the metastatic niche. Macrophage diversity and heterogeneity are well-established, with their phenotype and function tightly regulated by the local tissue environment ^75–77^.

Macrophages are known to promote bone metastasis, as broad macrophage targeting treatments like L-Clodrosome or CSF1R inhibition has shown efficacy in reducing bone metastasis outgrowth *in vivo* ^78^. However, the specific roles of macrophage subpopulations in supporting bone metastatic tumor outgrowth remains poorly defined CD204^high^IL4R^high^ macrophages derived from Ly6C^+^CCR2^+^ inflammatory monocytes have been shown to be important for bone metastasis outgrowth ^78^. Through the niche labeling, we found two macrophage clusters were enriched in the niche: Mac_c (*Vcam1*^+^*Cd163*^+^*Ccr3*^+^) and Mac_a (*Ccr2^+^Il4ra^+^*), which might correspond to CD204^high^IL4R^high^ macrophages. The niche-enriched *Vcam1*^+^*Cd163*^+^*Ccr3*^+^ macrophage subpopulation is distinct from the Cd204^high^Il4r^high^ population, indicating that different macrophage populations may play different roles and cooperate in supporting the metastatic growth of tumor cells in bone. The *Vcam1*^+^*Cd163*^+^*Ccr3*^+^ macrophages closely resemble EBI macrophages, as demonstrated by their high expression of genes involved in iron recycling and transport, adhesion molecules, and key transcription factors associated with EBI macrophages. In addition to their increased abundance in bone metastasis compared to normal bone, *Vcam1*^+^*Cd163*^+^*Ccr3*^+^ macrophages are recruited by metastatic tumors, relocating from the EBI niche to become highly enriched in the bone metastatic niche, where they facilitate tumor growth by rerouting iron recycling to benefit the tumors Emerging data suggest that tissue macrophages regulate iron availability in various environments, contributing to tissue homeostasis. For example, CD169^+^ macrophages control local iron availability, influencing HSC self-renewal and fate decisions under microbiota regulation ^80^. Microglia, brain resident macrophages, were found to play key roles in iron transport and metabolism in the central nervous system (CNS) ^81,82^. When cancer cells metastasize to the cerebrospinal fluid, they outcompete macrophages for iron by expressing iron-binding protein lipocalin-2 (LCN2) and its receptor SCL22A17 ^83^. Our study, along with these findings, underscore the critical role of macrophages in regulating local iron availability and supporting tumor growth. Furthermore, we discovered that tumor cells not only utilize iron for growth but use this resource to develop adaptive mechanisms to survive in the hypoxic bone environment. Tumor cells respond to increased iron flux by upregulating the heme synthesis while downregulating heme degradation pathway, enabling them to produce more heme. Simultaneously, they synthesize globin subunits for hemoglobin formation in response to hypoxia, thus displaying some of the hallmark cellular features of erythroblasts.

It has been increasingly recognized that tumor cells show phenotypic plasticity when they metastasize to distant organs ^35^. Previous research indicates that epithelial cells with mesenchymal traits can produce breast microcalcifications and develop osteoblast-like features in primary breast tumors ^84^. Clinically, mammographic detection of mammary microcalcifications is a key tool in breast cancer screening ^32^. Cancers that frequently metastasize to bone, such as breast, prostate, and lung cancers, exhibit an osteoblast-like phenotype through ectopic expression of osteoblast markers, bone matrix proteins and paracrine factors ^32–34,85^. These osteomimicry characteristics provided new strategies for predicting bone metastasis and combating multidrug resistance ^35^. In our study, we propose for the first time that metastatic bone tumors mimic another major bone cell type—erythroblasts, a phenomenon we refer to as “erythroblast mimicry”. First, tumor cells occupy the erythroblastic niche, utilizing iron-recycling macrophages to acquire iron for growth. Moreover, under hypoxic conditions, tumor cells upregulate *Gata1*, a key regulator for erythropoiesis, to induce hemoglobin gene expression to enhance their survival within the bone microenvironment. This research redefines the bone metastasis niche beyond traditional osteoclast-osteoblast interaction axis and reveals an important role of niche macrophages and its regulation of iron metabolism on bone metastasis development. Future studies should explore additional mechanisms involved in the recruitment of niche macrophage and development of erythroblast mimicry in metastatic tumors cells, as such studies will reveal opportunities for therapeutic intervention.

Anemia significantly affects both the quality of life and survival of cancer patients ^86–88^ and is the most common clinical presentation in breast cancer patients with bone metastasis ^89,90^. Furthermore, recent findings recent findings suggest that anemia negatively impacts responses to immune checkpoint inhibitors, where tumors convert erythroid precursors into myeloid cells, leading to a robust immunosuppressive phenotype ^91^. Understanding the mechanisms behind anemia and developing targeted treatments for cancer-associated anemia are crucial for improving not only quality of life but also survival and therapeutic response in cancer patients. Cancer-related anemia is believed to be caused by the activation of the inflammatory systems in cancer patients, leading to increased release of IL-6, TNFα, and IL-1. These pro-inflammatory cytokines stimulate hepcidin production in the liver, which negatively affects iron absorption and suppresses erythropoiesis in the bone marrow by limiting iron efflux from macrophages ^58^. In our study, we found that depletion of iron-rich macrophages with JEDI T cells prevents bone metastatic tumor growth and rescues the anemic phenotype, confirming the functional importance of these macrophages in cancer-associated anemia. We further discovered that bone metastatic tumor cells disrupt the normal distribution of iron-recycling macrophages in the bone marrow, leading to insufficient erythropoiesis. Tumors recruit *Vcam1*^+^*Cd163*^+^*Ccr3*^+^ macrophages from EBIs to the tumor edge, limiting the availability of iron for erythroblasts. This finding revealed the altered spatial distribution of EBI macrophages as a novel mechanism behind cancer-associated anemia. It also provides a potential explanation for refractory cancer-associated anemia, particularly in bone metastatic patients with reduced responses to erythropoietin (EPO) ^92^. Although anemia is widespread among cancer patients, its management remains controversial, with less than 50% of anemic cancer patients receiving treatment ^93,94^. Current therapies, including iron supplementation, erythropoiesis-stimulating agents, and RBC transfusion, have limitations due to safety concerns or potential adverse effect on stimulating tumor growth ^59,94^. Our research suggests potential new therapy strategies to treat or prevent anemia and reducing tumor growth in patients with bone metastasis by targeting the rerouting of EBI macrophages and restore their function in supporting erythropoiesis.

## Acknowledgments

This work was supported by the Charles H. Revson Senior Fellowship in Biomedical Science to Y. Han and grants from the Ludwig Foundation, Brewster Foundation, American Cancer Society, Breast Cancer Research Foundation and Susan G. Komen Foundation to Y. Kang, and R01CA183878, R01CA251950 and U01CA25355NIH to X. Zhang. We thank Y. Pritykin and T. Azad for their assistance with the single-cell RNA sequencing experiment and for their insightful discussions. We acknowledge the Princeton University Genomics Core Facility for library preparation and sequencing. We thank K. Rittenbach and the Molecular Biology Flow Cytometry Resource Facility, which is partially supported by the Rutgers Cancer Institute of New Jersey (NCI-CCSG P30CA072720-6852), for their assistance with cell sorting. We appreciate G. Laevsky and S. Wang at the Confocal Imaging Facility in the Department of Molecular Biology at Princeton University for assistance with confocal imaging. We thank J. Li at In Vivo Research Services (IVRS) Department of Rutgers University for mouse blood cell count analysis. We also thank Z. Cao for the assistance with cell culture experiments. We appreciate the entire Kang Lab for their insightful discussions throughout the project. We extend our gratitude to Dr. Daniel A. Notterman, Dr. Pengfei Zhang, Dr. Phoebe Carter and James Gow for their invaluable insights and discussion.

## Author Contributions

YH conceived the project, designed and performed the experiments, interpreted the results, analyzed the data, and co-wrote the manuscript. HS and SLD contributed to the single-cell RNA sequencing data analysis. ZX, FL, RLS and XZ provided the human bone metastasis single cell sequencing data. ZX also performed sectioning and staining of human samples. YW assisted with experiment design and data analysis. XH contributed to tissue processing and sectioning. KT assisted with animal experiments. WW and JM contributed to library preparation and sequencing. CD assisted with cell sorting. DB contributed to image quantification. YK conceived and supervised the project, designed the experiments, and co-wrote the manuscript.

## Methods

### Mice

All experimental protocols involving animals were carried out in accordance with the guidelines approved by the Institutional Animal Care and Use Committee (IACUC) of Princeton University. The facility maintained a controlled environment for the mice, which included temperatures of 20–22°C, a light cycle of 14 h:10 h, and a relative humidity of 40–70%. The mouse strains utilized, including C57BL/6J (RRID:IMSR_JAX: 000664), B6 albino (RRID:IMSR_JAX:000058), Spic-EGFP (RRID:IMSR_JAX:025673), C57BL/10 (RRID:IMSR_JAX:000666), JEDI (RRID:IMSR_JAX:028062) were obtained from The Jackson Laboratory. Mice genotyping was performed according to the JAX genotyping protocol. T cells isolated from C57BL/10 or JEDI mice were injected into F1 mice generated by crossing Spic-EGFP homozygous or heterozygous mice with C57BL/10 mice.

### Lentivirus production and infection

Plasmid pCDH-EF1a-eFFly-mCherry was a gift from Dr. Irmela Jeremias (Addgene, #104833), and pcPPT-mPGK-attR-sLP-mCherry-WPRE ^37,38^ was a gift from Dr. Ilaria Malanchi. Firefly luciferase-IRES-EGFP cassette was cloned into lentiviral vector pLEX under the control of the SFFV promoter. pLKO plasmids containing shRNA sequences targeting murine *Gata1*, along with two control shRNAs (Sigma, #shc001 and Sigma, #shc002v), were purchased from Sigma-Aldrich (St. Louis, MO, USA). The sequences of *Gata1* shRNAs are listed in Table S1. All plasmids were packaged into lentiviruses using HEK293T along with the helper plasmids VSVG and pAX2, following standard protocols. Lentiviruses harvested 48 hours post-transfection were used to infect target cells in the presence of 5 μg/ml polybrene.

The E0771 stable cell line was sequentially infected with Firefly luciferase-IRES-EGFP and sLP-mCherry-expressing viruses. EGFP and mCherry double-positive cells were isolated by FACS 48 hours post-infection with the sLP-mCherry virus. E0771, Py8119, and 4T1 cells were infected with eFFly-mCherry-expressing lentiviruses. mCherry- positive cells were selected using FACS.

### scRNA-seq sample preparation

E0771 breast cancer cells, engineered to express both EGFP and secreted mCherry, were injected into the caudal artery of 8- to 10-week-old female B6 albino mice to induce bone metastasis. Bone marrow cells were flushed from the femur and tibia using PBS containing 2% FBS. Red blood cells were eliminated using RBC lysis buffer. From the bone marrow of bone metastatic mice, niche cells (EGFP^-^mCherry^+^) and distal cells (EGFP^-^mCherry^-^) were isolated. Normal bone marrow cells (EGFP^-^mCherry^-^) were collected from healthy control mice.

Both bone metastatic tumors and primary tumors were dissociated using collagenase/hyaluronidase (Stemcell, #07912) and DNase I (Stemcell, #7900), processed with the gentleMACS™ Octo Dissociator using the 37C_m_TDK_1 program. Following RBC lysis (BioLegend, #420302), the cell suspension was filtered through a 70-μm strainer, stained with live dye (LIVE/DEAD™ Fixable Lime (506) Viability Kit, Invitrogen, #L3498) and antibodies, and prepared for FACS. For scRNA-seq library preparation, sorted cells from each sample were centrifuged and resuspended in PBS containing 0.5% BSA at 1,000 cells/μL.

### scRNA-seq library construction, sequencing and raw data processing

Single cell suspension samples were examined on Attune NxT Flow Cytometer for cell density and viability. Around 17,000 cells from each sample were loaded into each channel of Chip G on 10X Genomics Chromium X system using the Chromium Single Cell 3’ v3.1 Reagent Kits (10X Genomics Inc., CA) to capture individual cells and barcode their mRNA via reverse transcription following the manufacturer’s protocol. The cDNA samples were amplified and purified with Ampure XP magnetic beads (Beckman Coulter, CA), quantified by Qubit fluorometer (Invitrogen, CA), and examined on Bioanalyzer High Sensitivity DNA chips (Agilent, CA) for size distribution. Illumina sequencing libraries were generated from the amplified cDNA samples using the Illumina Tagment DNA Enzyme and Buffer kit (Illumina, CA). These libraries were examined by Qubit and Bioanalyzer, then pooled at equal molar amount and sequenced on Illumina NovaSeq 6000 S Prime flowcells as 28+94 nt pair-end reads following the standard Illumina protocol. Raw sequencing reads were filtered by Illumina NovaSeq Control Software and only the Pass-Filter (PF) reads were used for further analysis. After sample demultiplexing, raw PF reads were processed through the CellRanger software (10X Genomics Inc., CA, version 7.1.0) with genome reference data mm10-3.0 to obtain the gene expression levels in single cells and quality control matrices for each sample.

### Processing of scRNA-seq data

Cellranger processed count matrix data were loaded in R (version 4.3.1) using Seurat package (version 4.3.0) for further analysis. We first performed quality control and filtered out low-quality cells. Cells with fewer than 200 detected genes or more than 8,000 genes were excluded. Additionally, cells with more than 10% mitochondrial genes were filtered out. Doublets detected by the “scDblFinder” were also removed.

We normalized each dataset using the default parameters of the NormalizeData function. Following normalization, FindVariableFeatures function was employed to identify variable genes. For data integration, SelectIntegrationFeatures function was used with the top 5,000 variable genes to identify common genes used in the integration across all datasets. Subsequently, the FindIntegrationAnchors function was used with the “cca” method to get the integration anchors. These anchors were then used to integrate the data using the IntegrateData function with default settings.

### Unsupervised cell clustering

Unsupervised clustering of cells was performed with Seurat. We conducted ScaleData and RunPCA on the integrated data. The first 50 principal components were selected for clustering. Cell clusters were visualized using Uniform Manifold Approximation and Projection (UMAP) based on the top 30 principal components. The nearest neighbor graph was computed using the FindNeighbors function with the top 30 principal components. Unsupervised clustering was done using the FindClusters function with resolutions from 0.2 to 0.8. Based on previous knowledge and consistency within the different resolutions, we selected a resolution of 0.6 for three bone marrow samples and a resolution of 0.2 for the two tumor samples. Cluster marker genes were identified using the FindAllMarkers function by the Wilcoxon rank-sum test with the following parameters: only.pos = TRUE, min.pct = 0.25, and logfc.threshold = 0.25. Cell types were annotated using the Tabula Muris single-cell transcriptomic atlas as a reference ^41^. Gene signature scores were calculated using the AddModuleScore function. The gene expression levels shown in the manuscript were plotted using DotPlot, VlnPlot or FeaturePlot function within the RNA assay. Percentage stacked barplot was made with “ggplot2” (v3.5.1).

To identify differentially expressed genes (DEGs) in niche-specific population macrophages_c, we applied the FindMarkers function from Seurat to compare macrophages_c against macrophages_a and macrophages_b. Similarly, to identify DEGs in tumor cells, we used the FindMarkers function from Seurat to compare tumor clusters from bone metastatic sites and primary sites. Up-regulated genes (log2FC > 0.25, FDR-adjusted p-value < 0.05) were selected for enrichment/over-representation analysis using the enricher function from the clusterProfiler R package (v4.8.3) and the mouse genome annotation package org.Mm.eg.db (v3.17.0). The hallmark gene sets from the Molecular Signatures Database (MSigDB) were utilized for enrichment analysis through the msigdbr R package (v7.5.1). The top 10 enriched pathways, based on a p-adjusted method using Benjamini-Hochberg (pAdjustMethod = “BH”) with a p-value cutoff of 0.05, were visualized using the dotplot function.

### Bulk RNAseq sample preparation

Bone metastasis model was established by injecting E0771 breast cancer cells into the caudal artery of 8- to 10-week-old female B6 albino mice. Bone marrow cells were harvested by flushing the femur and tibia with PBS containing 2% FBS. Red blood cells were lysed using RBC lysis buffer (BioLegend, #420302), and the resulting cell suspension was filtered through a 70-μm strainer. Cells were then stained with LIVE/DEAD™ Fixable Lime (506) Viability Kit (Invitrogen, #L3498) and specific antibodies. General macrophages (CD11b^+^F4/80^+^) and niche-specific macrophages (VCAM1^+^CD163^+^CCR3^+^) were isolated from the bone marrow of both bone metastatic and healthy control mice via FACS. Total RNA was extracted using the RNeasy Micro Kit (Qiagen, #48500), following the manufacturer’s instructions.

### RNAseq library construction, sequencing, and data processing

The integrity of total RNA samples was assessed on the Bioanalyzer 2100 using RNA 6000 Pico chip (Agilent Technologies, CA). Messenger RNA was enriched from these samples using QIAseq FastSelect - rRNA HMR Kit (Qiagen, CA), fragmented and converted to cDNA then Illumina sequencing library using the PrepX RNA-seq library preparation protocol on the Apollo 324™ NGS Library Prep System (Takara Bio, CA).

Unique DNA barcode was incorporated into each library for sample demultiplexing. These RNA-seq libraries were examined on Agilent Bioanalyzer DNA High Sensitivity chips for size distribution, quantified by Qubit fluorometer (Invitrogen, CA), and pooled at equal molar amount. The library pools were denatured and sequenced on Illumina NovaSeq 6000 using the S Prime flowcells as pair-end 65 nt reads according to the manufacturer’s protocol. Raw sequencing reads were filtered by Illumina NovaSeq Control Software, only the Pass-Filter (PF) reads were de-multiplexed allowing 1 mismatch and used for further analysis.

Demultiplexed raw sequence files (FASTQ) were uploaded to the Partek® Flow® for analysis. Contaminant sequences (rDNA, mtrDNA, tRNA) were removed with Bowtie 2 - v2.2.5. Reads were aligned to mouse genome using STAR version 2.7.3a. Row gene counts were obtained by quantifying to mm10-Ensembl Transcripts release 100. Genes with less than 1 read were excluded.

Differential expression analysis of count data was performed using the DESeq2 R package (v1.40.2). The top 250 variable genes and differentially expressed genes were visualized using the pheatmap R package (v1.0.12). A pre-ranked list was generated based on the log2 fold change of all genes from VCAM1^+^CD163^+^CCR3^+^ cells compared with CD11b^+^F4/80^+^ macrophages from either normal or bone metastatic mice, using DESeq2 (v1.40.2). Low-expressed genes (normalized mean count <10) were filtered out before analysis. The ranked list was then used for GSEA ^95,96^, with the hallmark gene set from MSigDB using the fgsea R package (v1.26.0). The top 15 enriched pathways, ranked by adjusted p-value, were visualized using the ggplot2 R package (v3.5.1), with the Normalized Enrichment Score (NES) and FDR-adjusted p-value displayed. The enrichment plot for the Heme metabolism pathway was generated using the plotEnrichment function.

### Flow cytometry/FACS

Single cell suspension was prepared by filtering cells through a 70-μm strainer. Cells were then stained with the Live/Dead™ Fixable Aqua Dead Cell Stain Kit (Thermo Fisher, #L34966) on ice for 30 minutes, followed by a 5-minute incubation with Mouse BD Fc Block™ (BD Biosciences, #553142). Subsequently, cells were incubated with an antibody cocktail on ice for 45 minutes. Single color controls and fluorescence-minus-one (FMO) controls were prepared for gating. Compensation samples were created using the ArC™ Amine Reactive Compensation Bead Kit (Thermo Fisher, #A10346) for live dye stains and the Abc Total Antibody Compensation Bead Kit (Thermo Fisher, #A10497) for antibody stains.

Flow cytometry analysis was performed on Attune NxT Flow Cytometer (Thermo Fisher), and FACS was conducted on BD FACSAria Fusion Cell Sorter (BD Biosciences). Gating strategies are shown in Figure S2A, B. Flow data analysis was carried out using FlowJo software (FlowJo LLC). Statistical analysis and data visualization were performed with GraphPad Prism (GraphPad Software LLC).

Antibodies utilized for flow cytometry and FACS are detailed in Table S2.

### Prussian blue staining and quantification

Ferric iron stored in macrophages was detected using a Prussian blue staining kit (Abcam, #ab150674), following the manufacturer’s protocol. Briefly, working solution was prepared by mixing equal volumes of 20% hydrochloric acid and 10% potassium ferrocyanide solution immediately before use. Paraffin-embedded or frozen sections were rehydrated and immersed in this working solution for 20 minutes. Subsequently, slides were washed the three times in distilled water, counterstained with Nuclear Fast Red for 5 minutes, then rinsed twice in distilled water. Tissue sections are dehydrated sequentially in 95% and 100% alcohol, and then in xylene, with two changes for each. Finally, slides were covered by coverslip with a resinous mounting medium. Images were captured using the Aperio Versa 8 Slide scanner (Leica), with 20X objective lens under bright field.

Bright field images were then exported in TIFF format with Aperio ImageScope (Leica, v12.3.3) and uploaded in Image J (Fiji; National Institutes of Health, Bethesda, MD) for quantitative analysis. First, the input images were formatted as 24-bit RGB image. The Colour Deconvolution 2 ImageJ plugin was used to unmix the color into three channels: red (nuclear fast red stained all cells), blue (Prussian blue stained iron-rich macrophage) and brown (red blood cells). Region of interest (ROI) can be selected based on the tumor-stroma margin. Subsequent images underwent “Threshold” to label positively stained cells. The “Make Binary” and “Watershed” tools were used to better separate touching or overlapping objects. Signals of these positive stains were converted to digital signals using the “Analyze Particles” feature to obtain the X/Y coordinates, area and integrated intensity of each positive particles. This semi-automatic quantification method was verified by manual counts. Data visualization was done with R package ggplot2 and ggExtra, which adds marginal histograms/density plots to scatterplots/density plots.

### Immunofluorescent staining and quantification

For immunofluorescence, frozen sections were blocked using goat serum with 0.1% Triton-X100 at room temperature for one hour. The slides were then incubated with the primary antibody overnight at 4°C, followed by three washes with PBS. They were further incubated with a secondary antibody for one hour at room temperature, then washed three times with PBS. Before mounting with ProLong Gold antifade mountant (Molecular Probes, #P36930), all slides were incubated with DAPI (Thermo Fisher, #564907) for 5 minutes. Antibodies used for immunofluorescent staining are listed in Table S2. Images were captured using either the STELLARIS Confocal Microscope (Leica) or the Nikon A1 Confocal Microscope (Nikon) acquired at a resolution of 1,024 × 1,024 pixel.

Immunofluorescent images exported from Nikon NIS-Elements or Leica Application Suite X were uploaded in Image J (Fiji; National Institutes of Health, Bethesda, MD) for analysis. Images were split into individual channels and converted as 8-bit greyscale.

Fluorescent signals were converted to digital signals for data visualization, similar with Prussian blue stained images.

### RNA isolation and RT-qPCR

Total RNA samples were extracted from cells using the RNeasy Plus Mini Kit (Qiagen, #74136) according to the manufacturer’s instructions. One to two micrograms of RNA were reverse transcribed into cDNA using the SuperScript IV First-Strand Synthesis System (Thermo Fisher, #18091050) on a T100 Thermal Cycler (Bio-Rad). Quantitative PCR was conducted on the QuantStudio™ 3 Real-Time PCR System (Thermo Fisher) with PowerUp SYBR Green Master Mix (Applied Biosystems, #A25777). The sequences of the qPCR primers are listed in Table S3.

### Mouse blood cell count

Freshly collected mouse blood (30 μL) was transferred into an EDTA-anticoagulated collection tube (Greiner Bio-One, #450475). The samples were thoroughly mixed by inverting the tube 8 to 10 times. Blood cell count (CBC) analysis was then performed using the Element HT5 Auto Hematology Analyzer (Heska).

### Serum iron assay

Iron concentration and total iron-binding capacity (TIBC) in mouse serum were measured using the Total Iron-Binding Capacity and Serum Iron Assay ssKit (Abcam, #ab239715). For sample preparation, 10 μL of serum was added per well, and the volume was brought to 25 μL with TIBC Assay Buffer. For the Serum Iron Assay, after adding the buffer, the samples were incubated for 10 minutes at 37°C. For the TIBC Assay, iron solution was added to the buffer before incubation under the same conditions. Subsequently, 25 μL of TIBC Detector was added, followed by another 10- minute incubation at 37°C. Finally, TIBC Developer Solution was added and incubated for 10 minutes at 37°C. A standard curve was generated using 1 mM Iron Standard in each well to produce 0, 2, 4, 6, 8, and 10 nmol/well Iron Standard. Absorbance was measured at 570 nm for both standards and samples using the Varioskan LUX Multimode Microplate Reader (Thermo Fisher).

### Hemoglobin assay

Mouse hemoglobin levels were measured using the Hemoglobin Assay Kit (Abcam, #ab234046). Blood collected from the mouse tail was diluted 10-fold in ddH2O. Then, 10 μL of the diluted fresh blood was placed in each well of a 96-well clear plate with a flat bottom. Next, 90 μL of Hemoglobin Detector was added to all sample wells. The mixture was incubated at room temperature for 15 minutes. For the standard curve, 0, 5, 10, 15, and 20 μL of 1 g/dL Hemoglobin Standard were added to a series of wells in a 96-well and the volume in each was adjusted to 100 μL using Hemoglobin Detector, achieving final concentrations of 0, 50, 100, 150, and 200 mg/dL hemoglobin per well. The absorbance was measured at 575 nm using the Varioskan LUX Multimode Microplate Reader (Thermo Fisher). Hemoglobin concentration in the sample well was calculated using the standard curve after taking the dilution factor into account.

### T cell isolation

CD8^+^ T cells were isolated from the spleens of JEDI or C57BL/10 mice using the CD8a^+^ T Cell Isolation Kit (Miltenyi Biotec, #130–104-075). Briefly, 10^7^ spleen cells were resuspended in 40 μL of MACS buffer (PBS with 0.5% BSA and 2 mM EDTA). The cells were then incubated with 10 μL of a biotin-antibody cocktail at 4°C for 5 minutes. After this, 30 μL of MACS buffer and 20 μL of anti-biotin microbeads were added, and the mixture was incubated at 4°C for an additional 10 minutes. The cell suspension was then passed through an LS column. The flow-through that containing the enriched CD8+ T cells was collected for subsequent experiments

### Intra-cardiac (IC), caudal artery (CA) and tail vein injection

Female mice (8- to 12-week-old) were anesthetized with an intraperitoneal injection of ketamine and xylazine. For intra-cardiac (IC) injections, 50,000 tumor cells resuspended in 100 μL of PBS were administered slowly into the left ventricle using a 26-gauge needle. For caudal artery (CA) injections, 20,000-50,000 tumor cells in 50 μL of PBS were injected into the caudal artery in the tail using a 29-gauge needle. Successful IC and CA injections were confirmed by the rapid entrance of bright red blood in the needle hub. For tail vein injections, 1 million CD8^+^ T cells suspended in 100 μL of PBS were administered using a 29-gauge needle. For all injections, the needle was quickly withdrawn, and pressure was applied to minimize bleeding.

### Bioluminescence imaging (BLI)

Mice were anesthetized using Ketamine/Xylazine for *in vivo* bioluminescence imaging (BLI) with the IVIS Spectrum imaging system (PerkinElmer, USA). 100ul of 15mg/ml D-luciferin solution was injected to each mouse through the orbital plexus using an insulin needle. Images were acquired and quantified using Living Image 3D Analysis (PerkinElmer, USA).

### MicroCT scan

Mice were anesthetized using Ketamine/Xylazine for *in vivo* microCT scan with the Quantum GX2 microCT (PerkinElmer, USA). HU calibration and gain calibration were performed according to the manufacturer’s instructions prior to the scan. A large bore cover and small bed was used. Images were acquired at 90 kV and 88 uA, with a filter of 0.06 mm Cu and 0.5 mm Al, in high resolution mode for a 4-min scan. The Field of View (FOV) was 36 mm and the pixel size was 72 µm. Quantum GX2 software with a 3D viewer was used for 3D reconstruction and analysis.

### Cell culture

HEK293T cells (ATCC, #CRL-3216) were cultured in DMEM supplemented with 10% FBS, 20 mM HEPES, 100 IU/mL penicillin, 100 μg/mL streptomycin, and 10 mL/L amphotericin B. E0771 (a gift from Dr. Xiang H.-F. Zhang’s lab) and 4T1 cells (obtained from American Type Culture Collection) were cultured in RPMI-1640 media supplemented with 10% FBS, 20 mM HEPES, 100 IU/mL penicillin, 100 μg/mL streptomycin, and 10 mL/L amphotericin B. Py8119 (ATCC, CRL-3278), obtained from Dr. Weizhou Zhang’s lab, were cultured in Ham’s F12 media supplemented with 10% FBS, 20 ng/mL EGF, 5 μg/mL insulin, 2 μg/mL hydrocortisone, 100 IU/mL penicillin, 100 μg/mL streptomycin, and 10 mL/L amphotericin B.

### Macrophage culture

Bone marrow cells were collected as previously described and cultured in RPMI-1640 media supplemented with 10% FBS, 20 mM HEPES, 100 IU/mL penicillin, 100 μg/mL streptomycin, and 10 mL/L amphotericin B. A stock solution of heme (Hemin, Sigma, # 51280-1G) was prepared at 25 mg/mL (40 mM) in 0.15 M NaCl containing 10% NH4OH (Vehicle) as described in the literature^49^. Macrophages (Mac) were induced in the presence of 20 ng/ml M-CSF (PeproTech, # 315-02) plus vehicle. Iron rich macrophages (iMac) was induced in the presence of 20 ng/ml M-CSF plus 40 µM of Heme. Mac and iMac were collected 5 days after culture for subsequent tumor co-culture.

### Tumor-macrophage co-culture

Firefly luciferase-expressing E0771, 4T1 or Py8119 cells (2×10^4^ cells per well) and Mac or iMac (1×10^5^ cells per well) were seeded in the wells of 96-well plate. The co-cultured cells were treated with 1.5 µM of iron chelator Deferoxamine mesylate (DFO, Abcam, #ab120727), 150 µM of Ferroption (FPN) inhibitor VIT-2763/Vamifeport (Vam, MedChemExpress, #HY-112220), 10 µg/ml of iron-free apo-Transferrin (Trf, Sigma, #T1428). Tumor growth was assessed by measuring luciferase activity in the tumor cells, which stably express firefly luciferase. Luciferase activity was quantified by adding D-luciferin solution to a final concentration of 150 µg/ml, followed by measurement using a Promega GloMax Microplate Luminometer.

### Hypoxia culture

The medium containing HEPES to buffer pH was pre-equilibrated in a hypoxic chamber for 4-6 hours to remove oxygen. Cell cultures in the pre-equilibrated medium were placed into a Modular Incubator Chamber, where oxygen was displaced by infusing nitrogen (N2) to achieve hypoxic conditions with 2% O2 levels. A Petri dish containing sterile water was placed inside the chamber to ensure adequate humidification of the cultures. The Modular Incubator Chamber was returned to a conventional incubator for 24 or 48 hours. An identical culture was maintained in the same conventional incubator with 20% O2 to serve as a normoxia control.

### Cell viability assay

Cells were seeded in a 96-well plate and incubated under normoxic and hypoxic conditions for viability assays. The supernatant was centrifuged to collect suspended dead cells, and the attached cells were dissociated using trypsin. Suspended and attached cells were combined and stained for dead cells using the LIVE/DEAD™ Fixable Lime (506) Viability Kit (Invitrogen, #L3498) according to the manufacturer’s instructions, followed by analysis using the Attune NxT Flow Cytometer.

### Human bone metastasis samples

The collection of human bone metastasis samples was conducted in accordance with the Declaration of Helsinki and was approved by the Institutional Review Boards at Baylor College of Medicine (H-49396), The University of Texas MD Anderson Cancer Center (PA15-0225), and the University of Texas Medical Branch (H-46675). Written informed consent for the research use of their samples was obtained from all patients undergoing orthopedic surgery.

### Data availability

Raw and processed sequencing data have been deposited in the NCBI GEO database with accession number GSE270983 for mouse RNA-seq and scRNA-seq data and accession number GSE266330 for human scRNA-seq data ^74^. Data from the 615 patients in the Erasmus Medical Centre/Memorial Sloan-Kettering Cancer Center (EMC- MSK) cohort can be accessed with GEO Accession IDs GSE2603, GSE5327, GSE2034 and GSE122767 ^10^.

### Statistical Analysis

Statistical analyses were performed using GraphPad Prism (v10.3.0). Quantitative data are presented as mean ± SEM. Exact sample sizes, statistical test methods, and p-values are specified in the figure legends or figures. A significance level of p < 0.05 was considered statistically significant. GSEA was conducted using the R packages fgsea (v1.26.0) and clusterProfiler (v4.8.3). P-values were adjusted using the Benjamini-Hochberg method. Detailed information can be found in the Methods section. For non-quantitative data, including images from immunofluorescent staining, Prussian blue staining, and flow cytometry, findings were reproduced at least three times by replicating the experiments and/or using at least three biologically independent samples. Representative results are shown.

## SUPPLEMENTAL INFORMATION (Including Supplemental Figures S1-S7, Supplemental Tables S1-S3)

**Figure S1.**
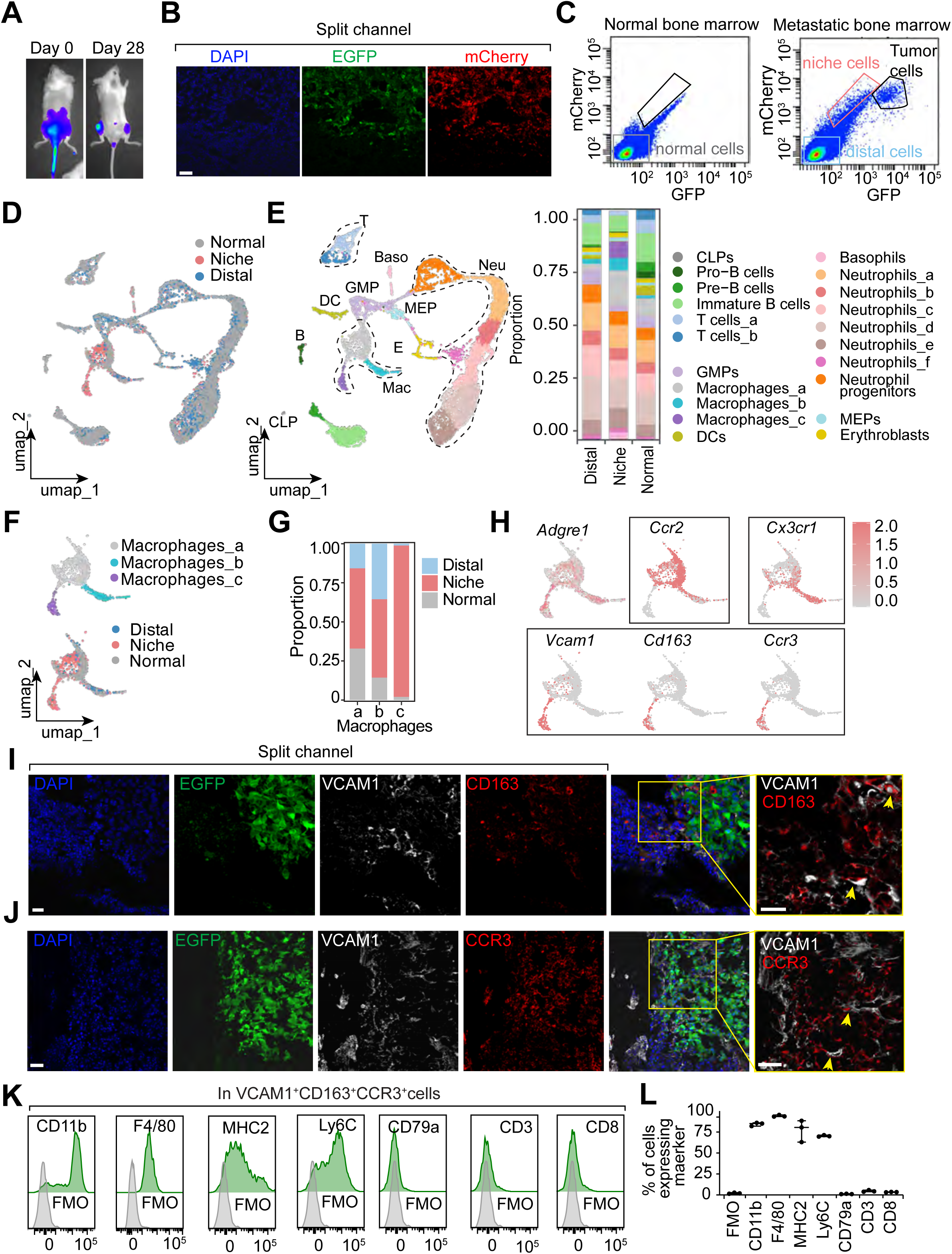
Discovery and validation of niche-specific macrophage in metastatic bone marrow. (**A**) Representative bioluminescence imaging (BLI) of sLP-mCherry/EGFP/luciferase-labeled E0771 cells at day 0 and day 28 following injection via caudal artery to 8- to 10- week-old female C57BL/6 albino mice. (**B**) Split channels of immunofluorescent images in Figure 1B. (**C**) Representative flow cytometry plots of normal (EGFP^-^mCherry^-^) bone marrow cells from healthy bone (pooled from three mice), niche (EGFP^-^mCherry^+^) and distal (EGFP^-^mCherry^-^) bone marrow cells from E0771 bone metastases (pooled from three mice). (**D**) Integrated UMAP plot of cells from normal, niche and distal bone marrow. (**E**) Integrated UMAP plot of cell clusters. Bar plot showing the proportion of each cell type in normal, niche and distal bone marrow. Colors indicate cell type. CLP: Common Lymphoid Progenitor, T: T cell, B: B cell, DC: Dendritic Cell, GMP: Granulocyte-Macrophage Progenitor, MEP: Megakaryocyte-Erythroid Progenitor, Ery: Erythroblast, Mac: Macrophage, Baso: Basophil, Neu: Neutrophil. (**F**) Integrated UMAP plots of three macrophage sub-clusters from normal, niche and distal bone marrow. (**G**) Bar plot showing the proportion of proportion of normal, niche and distal cells in each macrophage sub-cluster. (**H**) UMAP plots showing expression of *Adgre1* in all macrophages and enriched expression of *Ccr2* in Mac_a, *Cx3cr1* in Mac_b, and *Vcam1*, *Cd163* and *Ccr3* in Mac_c, colored according to z-score normalized expression levels. (**I**) Representative immunofluorescent images of DAPI (blue), EGFP (green), VCAM1 (white), and CD163 (red) in EGFP^+^ E0771 bone metastases. Scale bars, 50 μm. (**J**) Representative immunofluorescent images of DAPI (blue), EGFP (green), VCAM1 (white), and CCR3 (red) in EGFP^+^ E0771 bone metastases. Scale bars, 50 μm. (**K** and **L**) Representative flow cytometry histograms (K) and quantification (L) of myeloid and lymphoid marker expression in VCAM1^+^CD163^+^CCR3^+^ cells from E0771 bone metastases, with fluorescence-minus-one (FMO) as baseline control. n=3 biological replicates. Scale bars, 50 μm in (B), (I) and (J).

**Figure S2.**
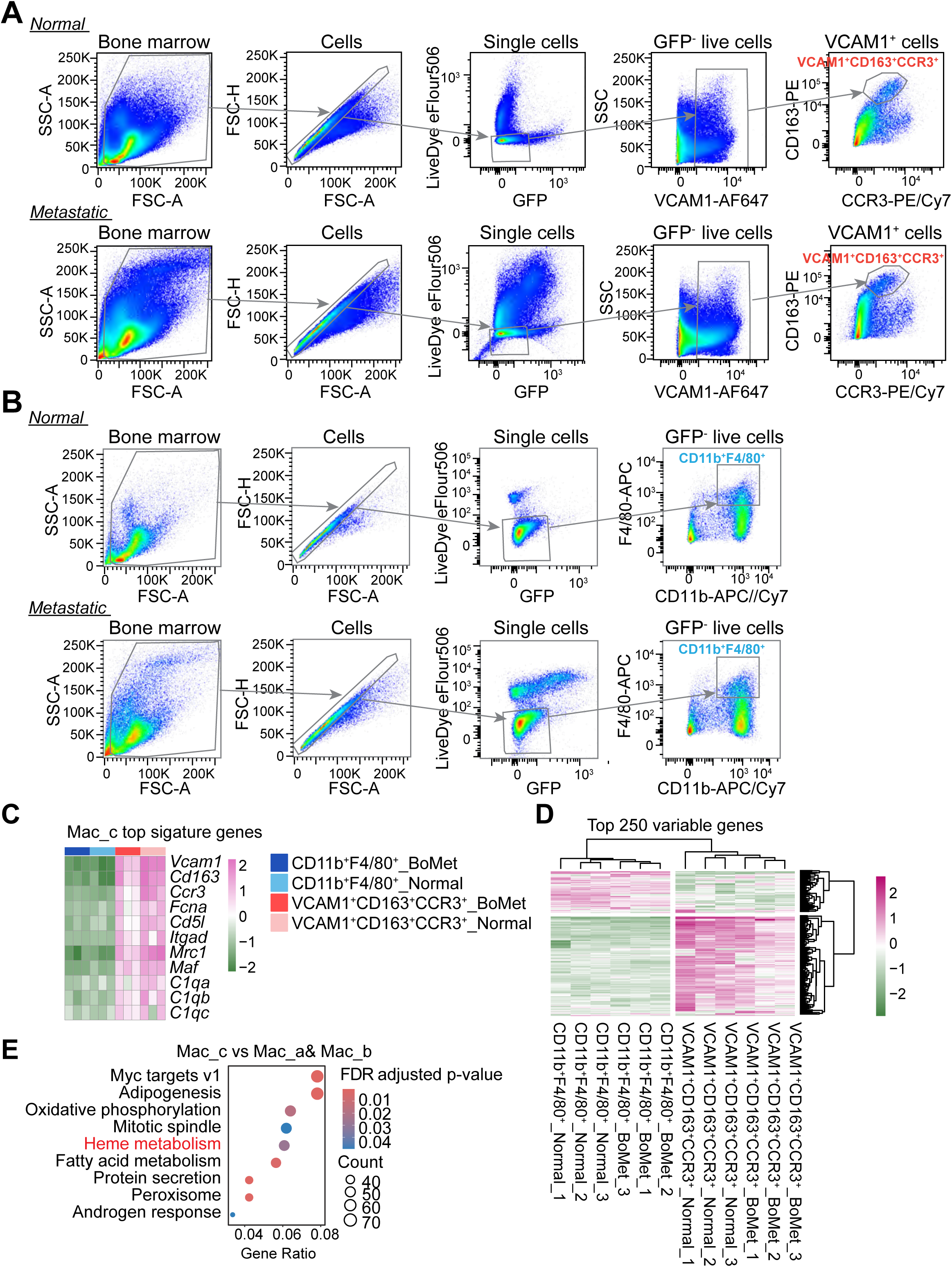
Sorting strategy and bulk RNA-seq analysis of niche-specific macrophages shown in Figure 2C to G. (**A** and **B**) Sorting strategy of VCAM1^+^CD163^+^CCR3^+^ (A) and CD11b^+^F4/80^+^ cells (B) from normal or metastatic bone marrow 28 days post-caudal artery injection of EGFP- labeled E0771 cells. (**C** and **D**) Heatmap showing the expression of Mac_c top signature genes from scRNA-seq data (C) and the top 250 variable genes (D) in the sorted VCAM1^+^CD163^+^CCR3^+^ and CD11b^+^F4/80^+^ cells, colored according to z-score normalized expression levels. (**E**) GSEA showing top differentially enriched hallmark pathways when comparing Mac_c with Mac_a and Mac_b in scRNA-seq data. Dot size represents the gene counts of enriched terms, colored according to FDR-adjusted p-value. The top 10 enriched pathways were identified using a p.adjust cutoff of 0.05 with the Benjamini-Hochberg method.

**Figure S3.**
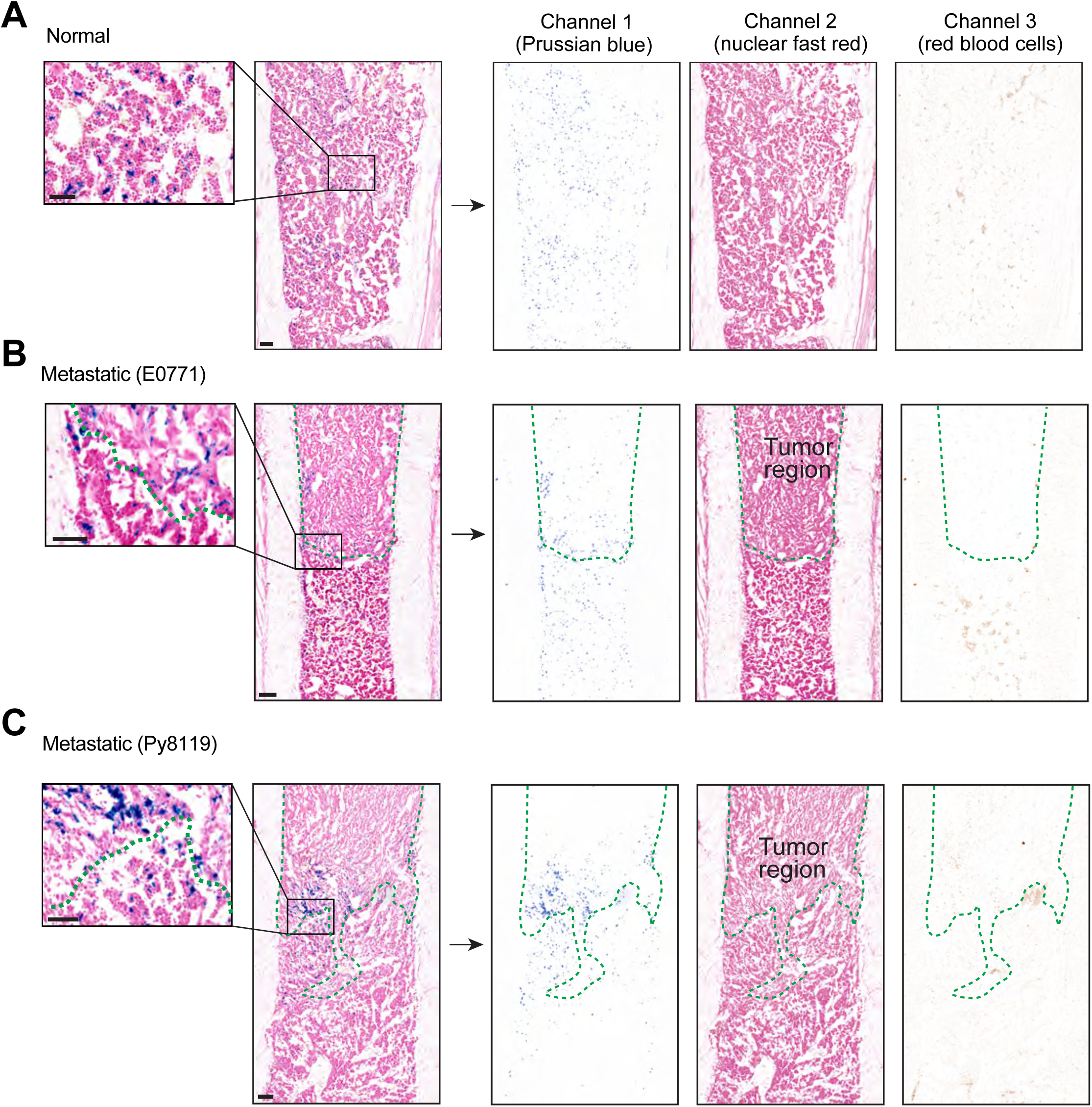
Prussian blue staining images used for quantification in Figure 3E. (**A**, **B** and **C**) Brightfield images of Prussian blue-stained normal bone section (A) and metastatic bone sections with E0771 (B) and Py8119 (C) tumors. Green dashed lines indicated tumor-stroma edge. Tumor regions were defined by distinct morphology and size of tumor cells compared to bone marrow cells. Immunofluorescence images of EGFP-expressing tumor cells in the adjacent slides were used as a reference. Prussian blue-stained images were split into three channels using Colour Deconvolution 2 ImageJ plugin: Channel 1 (Prussian blue stained iron-rich macrophage), Channel 2 (nuclear fast red stained all cells, tumor cells within the region of interest (ROI) were selected based on the tumor -stroma margin.), and Channel 3 (unstained brown red blood cells). Scale bars, 50 µm in the left panel and 100 µm in the right panel.

**Figure S4.**
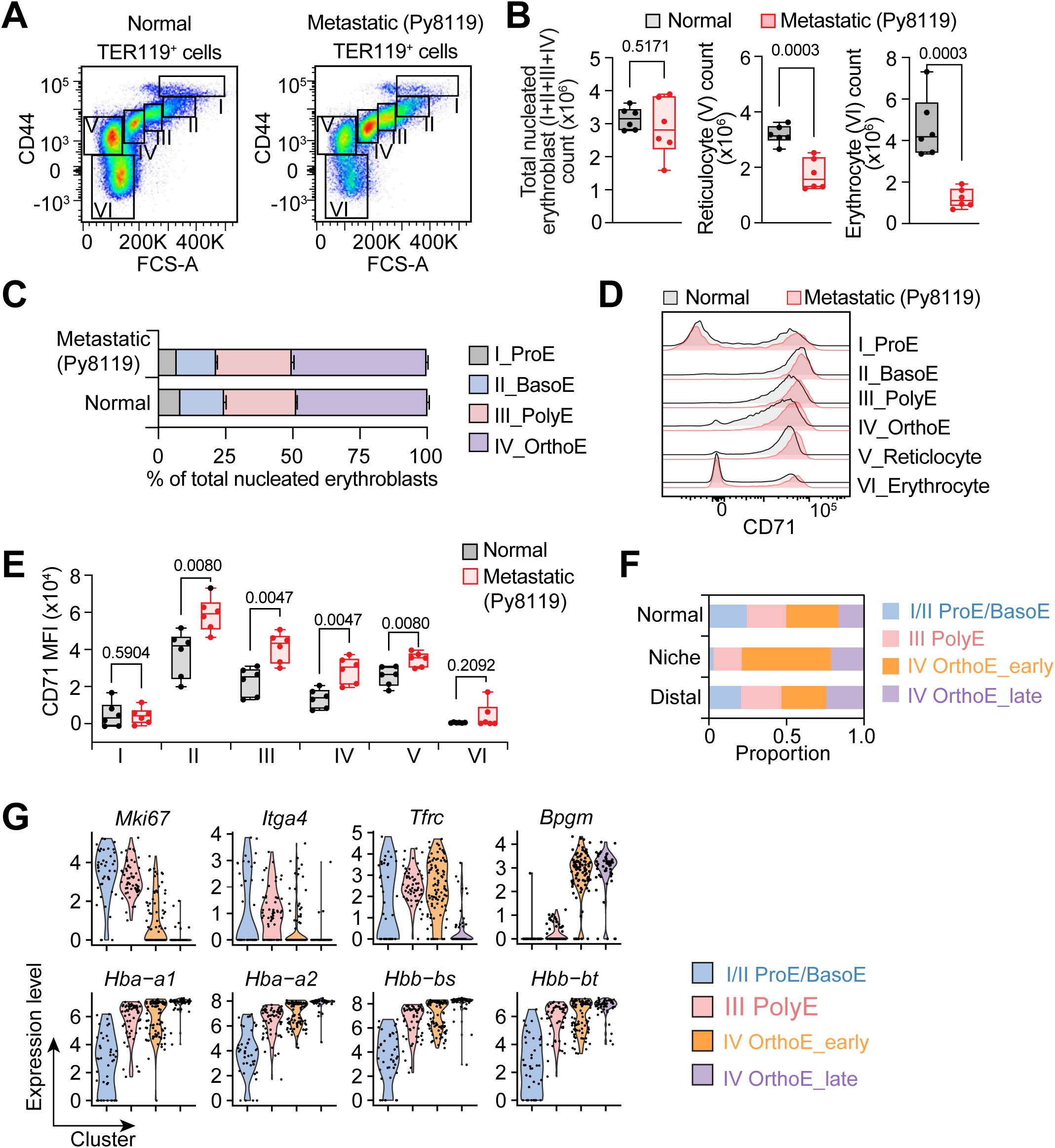
Inefficient erythropoiesis in the metastatic bone marrow. (**A**) Representative flow cytometry plots showing different stages of erythroid cells from normal and Py8119 metastatic bone marrow. (**B**) Quantification of the total nucleated erythroblast stages (I-IV), reticulocyte (V) and erythrocyte (VI) counts in normal or Py8119 metastatic bone marrow. n=6 biological replicates for each group. (**C**) Bar plots showing the proportions of ProE (I), BasoE (II), PolyE (III), and OrthoE (IV) within the total nucleated erythroblast population. (**D** and **E**) Representative flow cytometry histograms (D) and quantification of CD71 MFI (E) in different stages of erythroid cells from normal and Py8119 metastatic bone marrow. n=6 biological replicates for each group. (**F**) Bar plots showing the proportion of normal, niche and distal cells in each erythroblast sub-cluster. (**G**) Violin plots showing expression of erythroid differentiation genes in each erythroblast sub-cluster. All data is shown as mean ± SEM. Two-tailed unpaired Student’s t test (B), and multiple unpaired Student’s t test (E) were used to determine the statistical significance.

**Figure S5.**
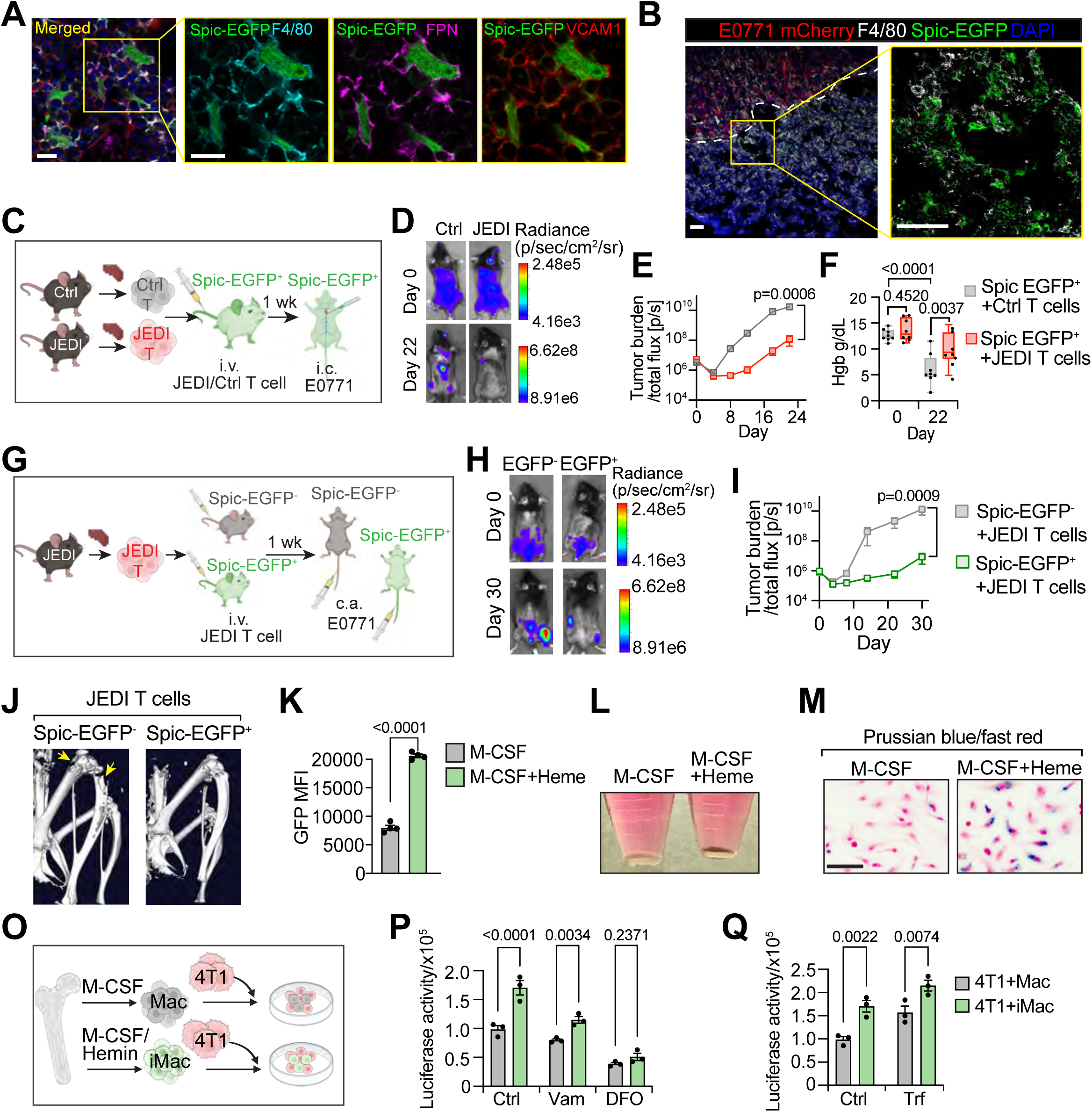
Iron-rich macrophages support tumor growth. (**A**) Representative immunofluorescent images showing expression of F4/80 (cyan), FPN (magenta), and VCAM1 (red), in Spic-EGFP (green) expressing cells in normal mouse bone marrow. (**B**) Representative immunofluorescent images showing expression of Spic-EGFP^+^ (green) F4/80^+^ (white) cells enriched at the edge of mCherry^+^ E0771 cells (red). The white dashed lines indicated tumor-stroma edge. (**C**) Schematic of the delivery of JEDI T cells or control T cells via intravenous injection (i.v.) one week prior to E0771 tumor cell injection through intracardiac injection (i.c.) into 8- to 12-week-old female Spic-EGFP^+^ mice. (**D**) Representative BLI images of Spic-EGFP^+^ mice injected with Ctrl or JEDI T cells, on day 0 and day 22 post-tumor inoculation (i.c.). (**E** and **F**) Tumor burden quantified by BLI (E) and HGB level (F) from day 0 to day 22 post-tumor inoculation (i.c.), in Spic-EGFP^+^ mice injected with Ctrl or JEDI T cells. n=8 replicates for each group. (**G**) Schematic of the delivery of JEDI T cells via intravenous injection (i.v.) one week prior to E0771 tumor cell injection through caudal artery (c.a.) injection into Spic-EGFP^+^ or Spic-EGFP^-^ mice. (**H**) Representative BLI images of Spic- EGFP^+^ mice injected with Ctrl or JEDI T cells, on day 0 and day 30 post-tumor inoculation (c.a.). (**I**) Tumor burden quantified by BLI from day 0 to day 30 post-tumor inoculation (c.a.), in Spic-EGFP^+^ or Spic-EGFP^-^ mice injected with JEDI T cells. n=5 replicates for Spic-EGFP^-^ group and n=9 replicates for Spic-EGFP^+^ group. (**J**) Representative microCT scans of bones from 8- to 12-week-old female Spic-EGFP^+^ or Spic-EGFP^-^ mice injected with JEDI T cells, collected on day 30 post-tumor inoculation. Bone metastases indicated by yellow arrow. (**K**) Quantification Spic-EGFP MFI in Mac and iMac by flow cytometry. n=3 biological replicates for each group. (**L**) Representative images of Mac or iMac pellets. iMac displayed a brown color after hemin uptake for 5 days. (**M**) Reprehensive bright-field images of Prussian blue stain showing iron storage in iMac, but not Mac. (**O**) Schematic of 4T1 co-culture with Mac or iMac. (**P** and **Q**) Luciferase activity showing 4T1 cell growth when co-cultured with Mac (n=3) or iMac, treated with iron chelator Deferoxamine (DFO) and the iron exporter FPN inhibitor vamifeport (Vam) (P) or treated with iron-binding protein transferrin (Q). n=3 replicates for each group. Scale bars, 20 µm in (A), 50 µm in (B and L). All data is shown as mean ± SEM. Two-tailed unpaired Student’s t test (M), and Two-Way ANOVA (E, F, P, Q) were used to determine the statistical significance.

**Figure S6.**
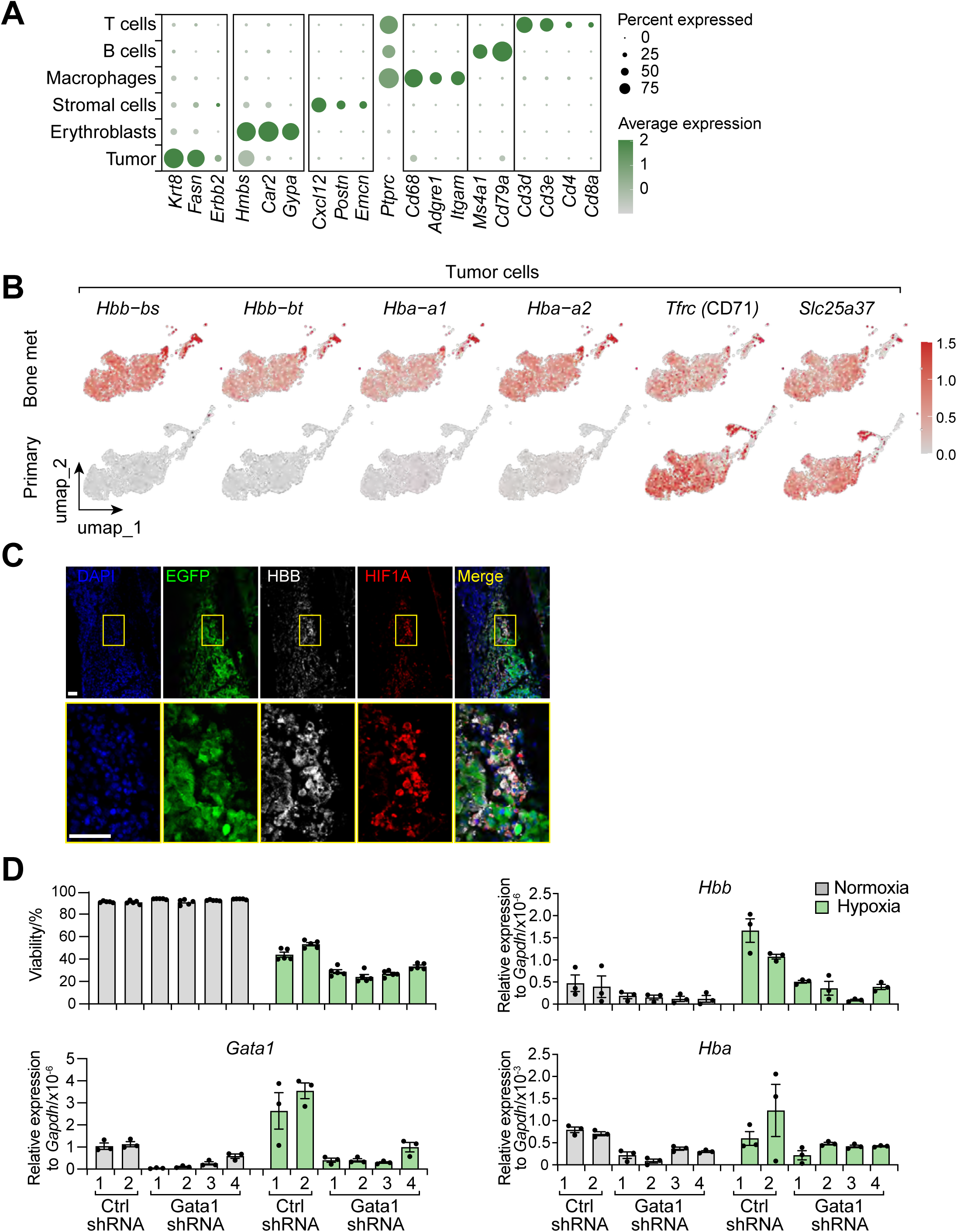
Bone metastatic tumor cells up-regulate globin gene expression in response to hypoxia. (**A**) Bubble plot showing marker genes of each cluster. Dot size indicates the fraction of expressing cells, colored according to z-score normalized expression levels. (**B**) UMAP plots showing expression of indicated genes in tumor cell cluster from bone metastases or primary site, colored according to z-score normalized expression levels. (**C**) Representative immunofluorescent images showing co-expression of HIF1A (red), and HBB (white) in EGFP^+^ E0771 cells (green). (**D**) Viability (48h after culture) and expression levels of *Gata1, Hbb* and *Hba* (24h after culture) in individual 2 control and 4 *Gata1* shRNA knockdown E0771 cell lines. n=3 replicates for each group. Scale bars, 50 µm in the left panel and 10 µm in the right panel. All data is shown as mean ± SEM.

**Figure S7.**
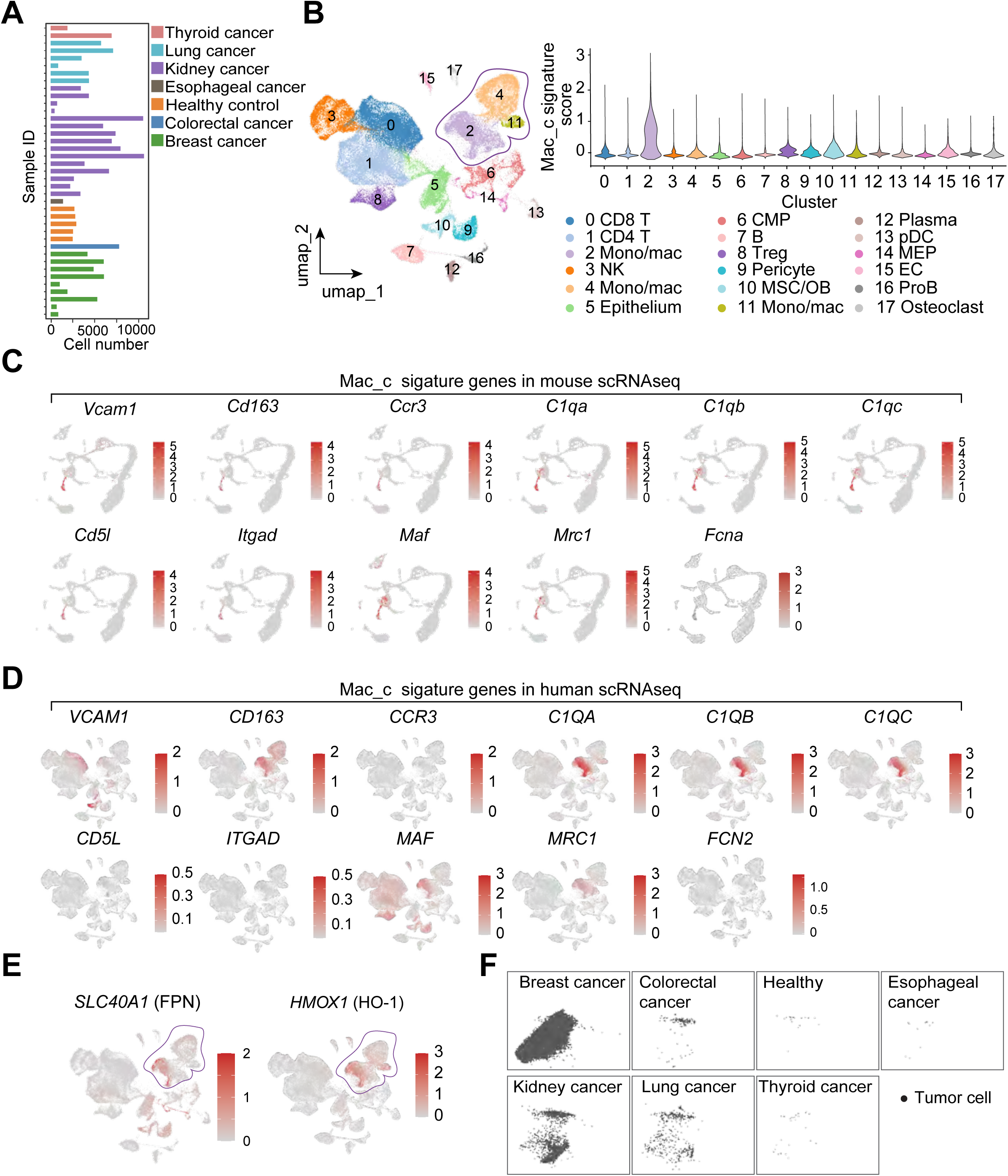
Validation of iron-rich macrophage in human bone metastasis samples. (**A**) Bar plot displaying the cell count analyzed for each sample, with bars colored according to cancer types. Sample collected from a total of 34 cancer patients (Breast cancer (n=9), colorectal cancer (n=1), esophageal cancer (n=1), kidney cancer (n=15), lung cancer (n=6), and thyroid cancer (n=2)) and 5 healthy donors. (**B**) UMAP plots of unbiased clustering of cells from human bone metastases from various cancer types following scRNA-seq analysis. Cell type annotation and detailed patient information can be found in the source study ^74^. Violin plots showing expression of the Mac_c signature score in different cell types. CMP: Common Myeloid Progenitor, OB: osteoblast, MSC: mesenchymal strom cells, Mac: Macrophage; pDC: Plasmacytoid Dendritic Cell, MEP: Megakaryocyte-Erythroid Progenitor, EC: Endothelial cell. OC: osteoclast. (**C** and **D**) Expression of individual Mac_c signature genes in the mouse bone marrow (C) and human bone metastases (D) data. Mac_c signature genes are also shown in Figure S2C. (**E**) UMAP plots showing *SLC40A1* (FPN) and *HMOX1* (HO-1) expression in all the cells. (**F**) UMAP plots showing epithelium (cancer cells) in different cancer types.

