## Supplemental Tables for "Niche Macrophages Recycle Iron to Tumor Cells and Foster Erythroblast Mimicry to Promote Bone Metastasis and Anemia"

**Supplementary Tables:**

**Table S1.** shRNA sequences used in the study.

**Table S2.** Antibodies used in this study with accompanying application and dilution notes.

**Table S3.** Quantitative RT-PCR primer sequences used in the study.

**Table S1. shRNA sequences used in the study.**

| Name | Source | Cat# | Target Sequence |
|---|---|---|---|
| GATA1 ShRNA_1 | Sigma | TRCN0000071609 | CCACTGACCATGAGGAAAGAT |
| GATA1 ShRNA_2 | Sigma | TRCN0000071610 | GTTTGGATGCAGCATCTTCTT |
| GATA1 ShRNA_3 | Sigma | TRCN0000071611 | CCTCTATTTCAAGCTCCATCA |
| GATA1 ShRNA_4 | Sigma | TRCN0000436155 | ACTGAGATTCAGGCATGTATT |

**Table S2. Antibodies used in this study with accompanying application and dilution notes**

| Target | Clone | Host | Source | Catalog | Application | Dilution |
|---|---|---|---|---|---|---|
| VCAM1-Alexa Fluor 647 | [illegible]29 (MVCAM.A) | | BioLegend | 105712 | Flow cytometry | 1:500 |
| CD163-PE | TNKUPJ | | Thermo Fisher Scientific | 12163180 | Flow cytometry | 1:200 |
| CCR3-PE/Cy7 | J073E5 | | BioLegend | 144513 | Flow cytometry | 1:200 |
| CD11b-APC-eFluor 780 | M1/70 | | Thermo Fisher Scientific | 47-0112-82 | Flow cytometry | 1:200 |
| F4/80-Super Bright 702 | BM8 | | Thermo Fisher Scientific | 67-4801-82 | Flow cytometry | 1:100 |
| MHC Class II-APC-eFluor 780 | M5/114.15.2 | | Thermo Fisher Scientific | 47-5321-82 | Flow cytometry | 1:200 |
| Ly-6C-APC/Cy7 | HK1.4 | | BioLegend | 128025 | Flow cytometry | 1:200 |
| Cd79a-APC-eFluor 780 | HM47 | | Thermo Fisher Scientific | 47-0792-42 | Flow cytometry | 1:200 |
| CD3-APC/Cy7 | 17A2 | | BioLegend | 100222 | Flow cytometry | 1:200 |
| CD8a-APC/Cy7 | 53-6.7 | | BioLegend | 100714 | Flow cytometry | 1:200 |
| CD45 PerCP-Cy5.5 | 30-F11 | | eBioscience | 45-0451-82 | Flow cytometry | 1:200 |
| Cd71-APC-eFluor 780 | RI7 217.1.4 | | Thermo Fisher Scientific | 47071182 | Flow cytometry | 1:200 |
| TER119-Super Bright™ 702 | TER-119 | | Thermo Fisher Scientific | 67-5921-82 | Flow cytometry | 1:200 |
| CD44-PE/Cy7 | IM7 | | BioLegend | 103030 | Flow cytometry | 1:200 |
| VCAM-1 | 1.4C3 | Mouse | Thermo Fisher Scientific | MA5-11447 | Immunofluorescence | 1:500 |
| CD163 | TNKUPJ | Rabbit | Thermo Fisher Scientific | 14-1631-82 | Immunofluorescence | 1:200 |
| CCR3 | - | Goat | Abcam | ab25789 | Immunofluorescence | 1:200 |
| F4/80 | D2S9R | Rabbit | Cell Signaling | 70076T | Immunofluorescence | 1:500 |
| F4/80 | BM8.1 | Rat | Cell Signaling | 71299S | Immunofluorescence | 1:500 |
| CD71 | H68.4 | Mouse | Thermo Fisher Scientific | 13-6800 | Immunofluorescence | 1:200 |
| Ferroportin/SLC40A1 | - | Rabbit | Novus Biologicals | NBP1-21502 | Immunofluorescence | 1:200 |
| HIF-1 alpha | mgc3 | Mouse | Abcam | ab16066 | Immunofluorescence | 1:200 |
| Hemoglobin alpha (HBA) | SN70-09 | Rabbit | Thermo Fisher Scientific | MA5-32328 | Immunofluorescence | 1:200 |
| Hemoglobin beta (HBB) | - | Rabbit | Thermo Fisher Scientific | PA5-60287 | Immunofluorescence | 1:200 |
| CD68 | - | Rabbit | Proteintech | 25747-1-AP | Immunofluorescence | 1:200 |
| Cytokeratin 8 | - | Rat | DSHB | TROMA-I | Immunofluorescence | 1:100 |
| Cytokeratin 19 | - | Rat | DSHB | TROMA-III | Immunofluorescence | 1:100 |
| anti-Rabbit Alexa Fluor 594 | - | Donkey | Thermo Fisher Scientific | A-21207 | Immunofluorescence | 1:1000 |
| anti-Rabbit Alexa Fluor 647 | - | Donkey | Thermo Fisher Scientific | A-31573 | Immunofluorescence | 1:1000 |
| anti-Mouse Alexa Fluor 647 | - | Goat | Thermo Fisher Scientific | A-21235 | Immunofluorescence | 1:1000 |
| anti Mouse Alexa Fluor 568 | - | Goat | Thermo Fisher Scientific | A-11031 | Immunofluorescence | 1:1000 |
| anti-Goat Alexa Fluor Plus 647 | - | Donkey | Thermo Fisher Scientific | A-32849 | Immunofluorescence | 1:1000 |
| anti-Goat Alexa Fluor 594 | - | Donkey | Thermo Fisher Scientific | A-11058 | Immunofluorescence | 1:1000 |
| anti-Rat Alexa Fluor 647 | - | Goat | Thermo Fisher Scientific | A-21247 | Immunofluorescence | 1:1000 |

**Table S3. Quantitative RT-PCR primer sequences used in the study.**

| Gene | Species | Direction | Sequences |
|---|---|---|---|
| *Gapdh* | Mouse | Forward | TCCCACTCTTCCACCTTCGATGC |
| | Mouse | Reverse | GGGTCTGGGATGGAAATTGTGAGG |
| *Hba* | Mouse | Forward | CACCACCAAGACCTACTTTCC |
| | Mouse | Reverse | CAGTGGCTCAGGAGCTTGA |
| *Hbb* | Mouse | Forward | GCACCTGACTGATGCTGAGAA |
| | Mouse | Reverse | ACTTCATCGGGGTTCACCTTT |
| *Klf1* | Mouse | Forward | CAGCTGAGACTGTCTTACCC |
| | Mouse | Reverse | AATCCTGCGTCTCCTCAGAC |
| *Runx1* | Mouse | Forward | GCCTCTCTGCAGAACTTTCC |
| | Mouse | Reverse | GACGGCAGAGTAGGGAACTG |
| *Gata1* | Mouse | Forward | AGGCCCTGGAAGACCAGGAAG |
| | Mouse | Reverse | AGAAAGGACTGGGAAAGTCAGC |
| *Gata2* | Mouse | Forward | CACCCCGCCGTATTGAATG |
| | Mouse | Reverse | CCTGCGAGTCGAGATGGTTG |
| *Klf4* | Mouse | Forward | GTGCCCCGACTAACCGTTG |
| | Mouse | Reverse | GTCGTTGAACTCCTCGGTCT |
| *Sox6* | Mouse | Forward | GGTCATGTTTCCCACCCACAA |
| | Mouse | Reverse | TTCAGAGGGGTCCAAATTCCT |
